# Optogenetic disruption of neural dynamics in the prefrontal cortex impaired spatial learning

**DOI:** 10.1101/2025.04.07.647582

**Authors:** Ignacio Negrón-Oyarzo, Tatiana Dib, Lorena Chacana-Véliz, Danae Barria, Koyam Morales-Weil, Marco Fuenzalida, Nelson Espinosa

## Abstract

1.

Cortical neural activity is highly dynamical at several temporal scales, property that has been postulated to be critical for the emergence of specific patterns supporting cognitive operations. During spatial learning, task-associated activity patterns gradually develop in the medial prefrontal cortex (mPFC) as the subject acquires experience. If neural activity dynamics is required in the mPFC for spatial learning is still unclear. Here we show that optogenetic entrainment of neural activity the mPFC disrupted local oscillatory and single-neuron dynamics. When applied during spatial training, optogenetic entrainment impaired behavioral performance and navigation strategy progression, a hallmark of spatial learning supported by the mPFC. Also, optogenetic entrainment blocked the emergence of learning-related activity patterns such as cross-frequency coupling and firing patterns signaling efficient goal approaching. Importantly, during memory retrieval, training-stimulated mice showed impaired performance in the absence of optogenetic stimulation. This evidence show that neural activity dynamics in the mPFC is crucial for spatial learning.

**Significance:** Cortical neural activity is highly fluctuating at several temporal scales, which has been postulated to be critical for cognitive functions. The medial prefrontal cortex (mPFC) is involved in the learning-related optimization of behavioral responses; however, it is unknown whether the dynamics of neural activity in the mPFC is required for learning. Here we found that optogenetic disruption of ongoing neural activity dynamics in the mPFC during training in a spatial memory acquisition task impaired spatial learning and hindered the emergence of learning-related activity patterns. These findings provide insight into the role of prefrontal activity dynamics in cognitive functions.

## 2.1. INTRODUCTION

Survival and well-being depend largely on the ability to incorporate new knowledge into future behavior. Thus, through learning, goal-oriented behavior is gradually optimized as the subject accumulates experience. This process is supported by the gradual experience-dependent development of neural activity patterns representing relevant neural operations for task achievement (Astorga et al., 2022; Benchenane et al., 2010; Hwang et al., 2019). It has been argued that the temporal variability of ongoing cortical activity (i.e. its dynamics) may be critical for the development of neural activity patterns supporting cognitive operations (Deco et al., 2011; Garrett et al., 2013; Stein et al., 2005; Waschke et al., 2021). The rationale is that neural dynamics emerges from the metastable transitions between synchronized and desynchronized states of neural populations (Chialvo, 2010; Friston, 1997; Garrett et al., 2013; Hancock et al., 2024; Harris & Thiele, 2011; Kopell et al., 2014; O’Byrne & Jerbi, 2022), and therefore, represents multiple spontaneous neural operations internally generated (Buzsáki, 2006; Buzsáki et al., 2014; Waschke et al., 2021). Thus, by means of external influence (i.e., experiences), these ongoing internal operations are adjusted and updated through the gradual integration of novel and relevant information collected through experience (Buzsáki, 2006; Buzsáki et al., 2014), facilitating the transient and flexible engagement of neural networks into optimal operations according to task demands (Stam, 2005; Waschke et al., 2021) which manifest as emergent neural activity patterns. However, despite the solid theoretical underpinning, experimental evidence testing this issue is scarce.

The medial prefrontal cortex (mPFC), a structure involved in the goal-directed accommodation of behavior (Miller, 2000; Miller & Cohen, 2001), contributes to the purposeful guidance of actions as a consequence of accumulated experience (Eichenbaum, 2017; Preston & Eichenbaum, 2013; Schlichting & Preston, 2015). In spatial learning tasks, the mPFC participates in the progressive implementation of increasingly efficient navigation strategies (de Bruin et al., 1994; Kesner et al., 1989; Kolb et al., 1994; Patai & Spiers, 2021), a hallmark of spatial learning (Harrison et al., 2006; Ruediger et al., 2012). It has been observed that specific activity patterns both in the local field potential (LFP) and single neuron levels gradually develop in the mPFC along spatial learning (Baeg et al., 2007; Benchenane et al., 2010; García et al., 2025; Negrón-Oyarzo et al., 2018). During spatial navigation, theta oscillation (6-12 Hz) is a main macroscopic activity pattern observed in the mPFC (Jones & Wilson, 2005; O’Neill et al., 2013; Sirota et al., 2008). Several features of prefrontal theta, such as power, phase, and its coordination with gamma oscillations, emerge transiently at specific moments of cognitive relevance as long as the animal acquires experience, which is associated with successful learning (Benchenane et al., 2010; García et al., 2025; Negrón-Oyarzo et al., 2018; Tavares & Tort, 2022). Also, prefrontal theta coordinates local neuronal spiking, promoting the learning-associated emergence of firing patterns representing cognitively relevant operations, such as decision-making, strategy switching, or efficient strategy implementation (Benchenane et al., 2010; Negrón-Oyarzo et al., 2018).

Here, we hypothesize that neural activity dynamics is required for the development of learning-associated activity in the mPFC, and consequently, spatial learning. To test this possibility, we perturbed neural activity dynamics but maintained theta activity through optogenetic theta-frequency entrainment of prefrontal activity specifically during spatial training. Given that excitatory layer 5 pyramidal neurons are the main contributor to cortical LFP (Beltramo et al., 2013), which represent relevant features of the task (Negrón-Oyarzo et al., 2018), the specific cell-identity expression of opsins in those neurons allows for optogenetically entraining neuronal and oscillatory prefrontal activity. Thus, in combination with simultaneous neural activity recording and linear and non-linear measurement of neural activity dynamics (Waschke et al., 2021), we were able to establish the relationship between neural activity dynamics in the mPFC and spatial learning. Our results showed that optogenetic entrainment transiently disrupted neural dynamics. When applied during training, optogenetic entrainment impaired behavioral performance and navigation strategy progression, and blocked the emergence of learning-related activity patterns such as cross-frequency coupling and firing patterns signaling goal approaching. Finally we found that mice exposed to optogenetic entrainment during training showed impaired performance during memory retrieval. These results reveal the relevance of neural activity dynamics in the mPFC for spatial learning.

## 3. RESULTS

### 3.1. Optogenetic stimulation of layer 5 pyramidal neurons in the mPFC entrains single units and LFP and disturbs local neural activity dynamics in ChR2-expressing mice

To entrain neural activity in the mPFC we used the *Thy1-ChR2-EYFP* transgenic mice, which express ChR2 specifically in layer 5 pyramidal neurons in several cortical areas (Arenkiel et al., 2007), including the mPFC (**Fig. 1a**). We evaluated the minimum light intensity (λ = 473 nm) necessary to evoke action potentials. To this aim, whole-cell patch clamp recordings (**suppl. Fig. 1a**) were performed in slices containing the mPFC (WT; n = 5 neurons; ChR2; n = 6 neurons). The stimulation protocol consisted of pulses of 15 ms delivered at 10 Hz at several light power intensities from 0 to 1.5 mW (**Fig. 1b**), yielding a theoretical irradiance between 0.0 and 11.81 mW/mm^2^ (Deisseroth lab, n.d.). Stimulation at light intensity above 0.5 mW (irradiance = 3.94 mW/mm^2^) evoked action potentials exclusively in pyramidal neurons of layer 5 (**suppl. Fig. 1b**). No action potentials were observed at all tested light irradiances in layer 2/3 pyramidal neurons and interneurons in ChR2 mice, as well as in layer 2/3 and 5 pyramidal neurons in WT mice (**suppl. Fig. 1b**). We next tested the minimum light intensity to entrain prefrontal unitary activity *in vivo*. To this aim, WT and ChR2 mice were chronically implanted with opto-multielectrode arrays (Opto-MEA; **suppl. Fig. 2a-c**) and subjected to testing sessions in an open field (**Fig. 1c**), in which we evaluated the effect of different light intensities in the prefrontal neural activity. Each session was subdivided into three stages of 60 sec each: pre, stimulation, and post (**Fig. 1c**). Single-neuron and LFP activity were recorded in the three stages, whereas light stimulation (the same protocol used in patch clamp experiments; **Fig. 1b**) was delivered in the stimulation stage. Fiber optics was correctly positioned in the mPFC, as shown in **Fig. 1d**. No difference in basal firing rate was found between groups (P = 0.763; t-test; WT, n = 11 neurons; ChR2, n = 19 neurons; **suppl. Fig. 2d, e**). Light stimulation induced firing of prefrontal units at intensity above 0.5 mW, but significant differences were obtained at 1.0 mW (n = 25 units from 9 animals; effect of stimulation; P < 0.0001; effect of light intensity: P = 0.0058; **Fig. 1e, f** and **suppl. Fig. 2f**). No increased firing was detected at these intensities in WT mice (n = 19 units from 7 animals; effect of light intensity: P = 0.535; two-way ANOVA; **Fig. 1e, f** and **suppl. Fig. 2f**). Stimulation resulted in a stereotyped firing in ChR2 mice, as revealed by the distribution in the inter-spike intervals (ISI), an effect not observed in WT mice (**suppl. Fig. 2g**). Therefore, 1.0 mW was the minimal light intensity to entrain firing in the mPFC. At a macroscopic level, light stimulation produced an evident inflection in the amplitude of the LFP in ChR2 mice, but not in WT mice (**suppl. Fig. 3a**). LFP recordings and power spectral analysis revealed that theta-frequency light stimulation induced an evident increase in the 10 Hz power in ChR2 mice, but not in WT mice, which returned to basal levels in the post-stimulation period (WT: n = 8; ChR2: n = 9; effect of stimulation: P = 0.026, interaction between stimulation and group: P = 0.028; two-way ANOVA; *post hoc* comparison WT vs ChR2: P < 0.01; **Fig. 1g, h)**. This stimulation generated periodical harmonics every 10 Hz, a distinctive feature of pulsatile light simulation (**Fig. 1h**) (Ahlgrim & Manns, 2019; Bitzenhofer et al., 2017). No effect was observed for low and high-gamma frequency bands (effect of stimulation; low-gamma: P = 0.171; high-gamma: P = 0.169; two-way ANOVA; **suppl. Fig. 3b, c**).

**Figure 1.**
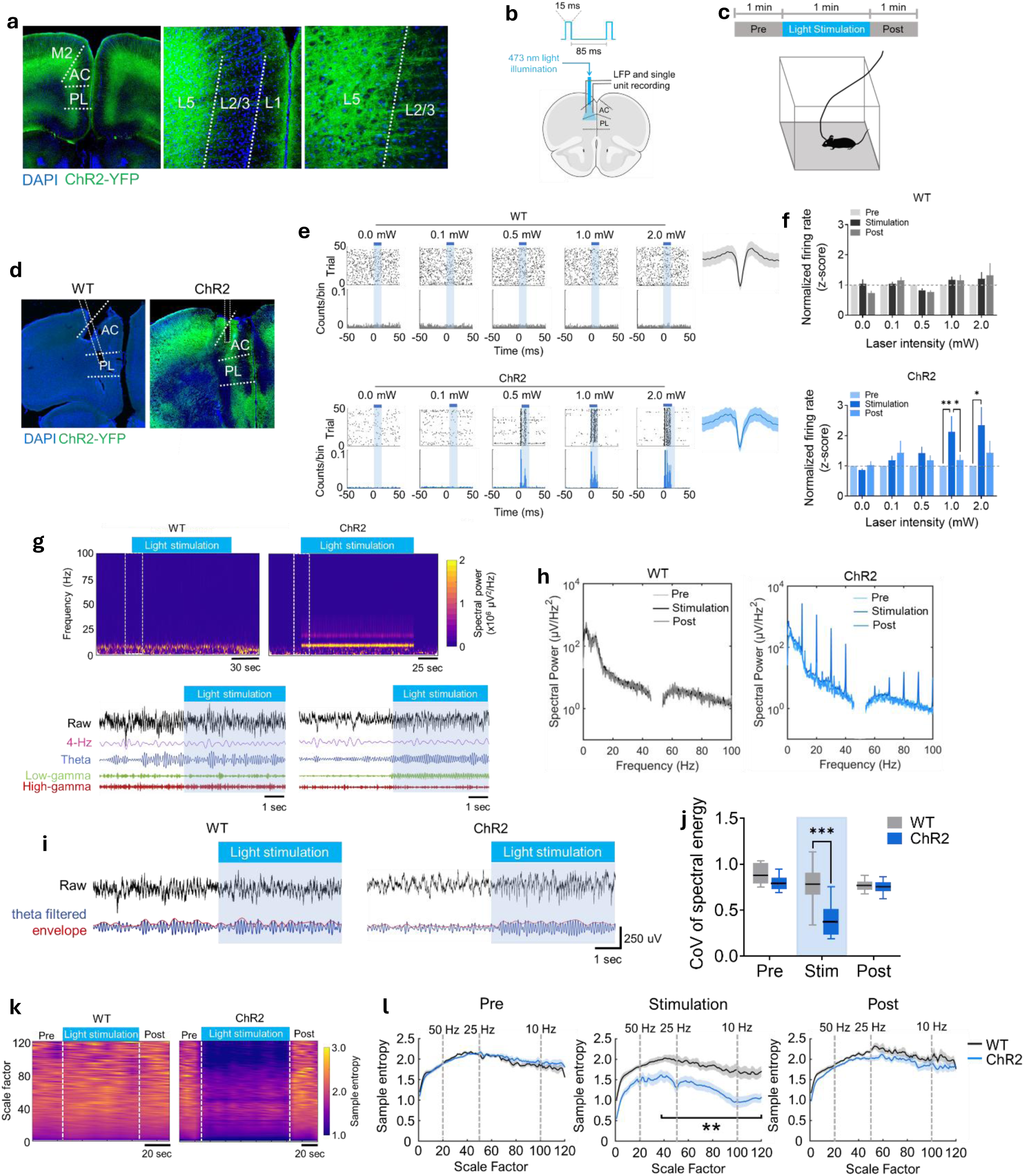
Optogenetic stimulation of layer 5 pyramidal neurons in the mPFC entrained single units and LFP and disturbed neural activity dynamics in ChR2-expressing mice. **a)** Confocal microphotography (magnification left, 4x; middle, 40x; right, 100x) showing the expression of ChR2-YFP protein in the mPFC of ChR2 (*Thy1-ChR2-YFP*) mice. M2: supplementary motor cortex; AC: anterior cingulate cortex; PL: prelimbic cortex; L1: layer 1; L2/3: layer 2/3; L5: layer 5. **b)** Schematic diagram of the experimental setup. WT and ChR2 mice were chronically implanted with a fiber optic surrounded by microwire electrodes for LFP and single unit recording in the mPFC. Blue light (473 nm) pulses (15 ms) at 10 Hz (inter-pulse duration: 85 ms) were delivered to both groups. c) Schematic diagram of the testing experiment: chronically opto-MEA implanted mice of both WT and ChR2 groups were subjected to simultaneous LFP and single unit recording in an open field during the pre, stimulation and post stages. Light stimulation was delivered at different light-power intensities from 0 to 2.0 mW specifically during the stimulation stage. **d)** Example of confocal microphotography showing the position of fiber optic in the mPFC of a WT (upper) and ChR2 mice (lower). AC: anterior cingulate cortex; PL: prelimbic cortex. **e)** Raster plot and peri-stimulus time histogram of examples of single units recorded in the mPFC of WT (upper) and ChR2 (lower) mice stimulated at different light-power intensities from 0 to 2.0 mW. The average waveform of the recorded unit is shown on the right. **f)** Bar chart comparing the mean normalized (z-scored) firing rate of all single units recorded from the mPFC of WT (upper) and ChR2 (lower) mice before (pre) during (stimulation) and after (post) light stimulation at different light-power intensities (from 0 to 2.0 mW). Data are presented as mean ± SEM. *: P < 0.05; **P < 0.01; Sidak’s multiple comparisons test after two-way ANOVA. **g)** Example of time-frequency color-coded analysis of spectral power (spectrogram; upper panel) and raw and filtered (4-Hz, theta, low-gamma, and high-gamma) LFP recording (lower panel) from the mPFC of WT (left) and ChR2 (right) mice during 10-Hz light stimulation. **h)** Average power spectral density analysis of the LFP recorded from the mPFC in WT and ChR2 mice during pre, stimulation and post stages. Data are presented as mean (solid line) ± SEM (shaded area). **i)** Example of raw (black) and theta-filtered (blue) and its envelope (red) of LFP recording from the mPFC of WT (upper) and ChR2 mice (lower) during light stimulation. **j)** Box plot comparing the CoV of spectral energy of theta frequency oscillations in the mPFC before (pre) during (stimulation) and after (post) light illumination in WT and ChR2 mice. The middle, bottom, and top lines of the box plot correspond to the median, lower, and upper quartiles, and the edges of the lower and upper whiskers correspond to the 5th and 95^th^ percentiles. ***: P < 0.001; Sidak’s multiple comparisons test after two-way ANOVA. **k)** Example of time-scale analysis of multiscale entropy (MSE) of LFP signal recorded from the mPFC of WT (left) and ChR2 (right) mice during 10-Hz light stimulation. l) Comparison of the average MSE of the LFP signals recorded from the mPFC of WT and ChR2 mice before (pre) during (stimulation) and after (post) 10-Hz light illumination. **: P < 0.01; Wilcoxon-signed rank test. Data are presented as mean (solid line) ± SEM (shaded area).

Next we evaluated the effect of entrainment on neural dynamics, for which we measured variance-based parameters, as the coefficient of variation (CoV) of spectral energy (Waschke et al., 2021). We found a significant decrease in CoV of spectral energy in the theta band in ChR2 mice specifically during stimulation (effect of group: P < 0.0001; effect of stimulation: P < 0.0001; interaction between group and stimulation: P < 0.0001; *post hoc* analysis WT vs ChR2 during stimulation: P < 0.0001; two-way ANOVA; **Fig. 1i, j**). We then evaluated neural dynamics through an entropy-based measurement (Waschke et al., 2021) as multi-scale entropy (MSE), a non-linear statistic that estimates the complexity of time series across multiple time scales (Costa et al., 2005). This method calculates entropy values on coarse-grained time series (fast to slow) and, therefore, allows the calculation of entropy in the frequency domain. We found a significant decrease in MSE during stimulation at a factor scale higher than 40 (oscillatory frequency under 25 Hz) in ChR2 mice compared to WT (P < 0.01; Wilcoxon rank-sum test; **Fig. 1k, l**). No differences were detected between groups in the pre- and post-stimulation stages at all scale factors (P > 0.05; Wilcoxon rank-sum test; **Fig. 1k, l**). We then evaluated whether prefrontal stimulation affected oscillatory dynamics in areas connected to the mPFC, such as the hippocampus (HPC) (Jay & Witter, 1991; Swanson, 1981). We found that the entrainment of prefrontal activity did not affect the spectral power of hippocampal theta oscillation (P = 0.934; one-way ANOVA), the CoV of spectral energy (P = 0.463; one-way ANOVA**; suppl. Fig. 4a-f**), and MSE (P > 0.05; Wilcoxon rank-sum test**; suppl. Fig. 4g, h)**. Together, these data suggest that entrainment of neural activity in the mPFC disrupts specifically the local neural dynamics.

### 3.2. Entrainment of neural activity in the mPFC during training impaired spatial learning

We then evaluated the effect of the entrainment of prefrontal activity on spatial learning. To this aim, we used the Barnes maze, a behavioral paradigm in which animals learn, through a series of exploratory trials, the spatial location of an escape from an aversive arena (schema in **Fig. 2a**). Each trial session is composed of three stages: start, navigation, and goal (**Fig. 2b**). WT and ChR2 opto-MEA implanted mice (WT: n = 7; ChR2: n = 9) were subjected to 4 training trials during 4 consecutive days, in which theta-frequency light stimulation was delivered exclusively during the navigation stage on all training sessions (**Fig. 2b**). WT-stimulated mice showed a tendency to navigate close to the escape throughout training, a typical hallmark of spatial learning **(Fig. 2c; suppl. video 1)**. This learning was manifested as the progressive reduction of escape latency, error counts (nose-poke in non-escape holes), and travelled distance to escape (**Fig. 2d** and **suppl. Fig. 5a, b**). Contrarily, ChR2-stimulated mice did not manifest changes in performance measurements, which remained significantly higher compared to WT mice (effect of group on escape latency, error counts, and distance travelled to escape: P < 0.000; two-way ANOVA; **Fig. 2d-e**, **suppl. Fig. 5a, b; suppl. video 2**). WT animals also displayed a decrease of escape nose-poke latency through training days (**Fig. 2e**), a more reliable indicator of behavioral performance in the Barnes maze (Harrison et al., 2006), together with significant changes in the cumulative distribution escape nose-poke latency between training days 1 and 4 (P = 0.046; Kolmogorov-Smirnov test; **suppl. Fig. 2c**). Contrarily, ChR2-mice displayed no changes in the escape nose-poke latency between training days 1 and 4 (P = 0.701; Kolmogorov-Smirnov test; **suppl. Fig. 5c**). Comparison between WT and ChR2 revealed significant differences in the escape nose-poke latency and the number of errors before escape nose-poke (P = 0.0015 and P = 0.013, respectively; two-way ANOVA; **Fig. 2e** and **suppl. Fig. 5d**). Consistent with behavioral performance, WT mice progressively increased their path efficiency and decreased their cumulative integrated path length (CIPL) to escape nose-poke across training days (**Fig. 2f, g**). Conversely, ChR2 mice showed no improvement in both parameters (**Fig. 2f, g**), and we found a significant effect of group in path efficiency and CIPL (P = 0.018 and P = 0.0002, respectively; two-way ANOVA; **Fig. 2f, g**). These differences were not attributable to locomotor activity, as we found no differences in mean and maximum speed (effect of group in mean speed: P = 0.194; effect of group in max. speed: P = 0.224, two-way ANOVA**; suppl. Fig. 5e, f**), or to the motivation to find the escape, as we found no differences in error incidence (errors per second; effect of group: P = 0.794; two-way ANOVA**; suppl. Fig. 5g**). Also, these behavioral differences were not attributable to genotype, as we found no differences between WT and ChR2-mice not subjected to light stimulation in escape nose-poke latency (P = 0.255), and errors (P = 0.160), path efficiency (P = 0.114), and CIPL to escape nose-poke (P = 0.831, two-way ANOVA**; suppl. Fig. 5h-l**). We then asked if prefrontal entrainment modified strategy progression. To this aim, we qualitatively classified the path to the escape into random, serial, and direct strategies (**Fig. 2h; suppl. video 1 and 2**). Strategies impact behavioral performance, as escape nose-poke latency, path efficiency, and CIPL were significantly affected by navigation strategies (P < 0.0001, one-way ANOVA; **suppl. Fig. 5m**). We found that WT mice significantly decreased the use of random and increased the use of direct strategy between the training days 1 and 4 (P = 0.045 and P = 0.004, respectively; Sidak’s multiple comparisons test after two-way ANOVA; **Fig. 2i, j)**. In contrast, ChR2 mice did not change the utilization of random and direct strategies between training days 1 and 4 (P = 0.627; P = 0.173, respectively; Sidak’s multiple comparisons test after two-way ANOVA; **Fig. 2i, j**). Indeed, we found significant differences between WT and ChR2 in the utilization of random and direct strategies on day 4 (random: P = 0.044; direct: P = 0.018; Sidak’s multiple comparisons test after two-way ANOVA) but not on day 1 (random: P = 0.217; direct: P = 0.990; Sidak’s multiple comparisons test after two-way ANOVA). These effects were not due to genotype, as we found in both WT and ChR2 mice significant differences between day 1 and 4 in the use of random (WT: P = 0.008; ChR2: P = 0.017; Sidak’s multiple comparisons test after two-way ANOVA) and direct strategies (WT: P = 0.002; ChR2: P = 0.001; Sidak’s multiple comparisons test after two-way ANOVA; **suppl. Fig. 5o**). Altogether, our behavioral results suggest that optogenetic entrainment of prefrontal activity impaired spatial learning in ChR2 mice, which was associated with a deficit in strategy progression across training.

**Figure 2.**
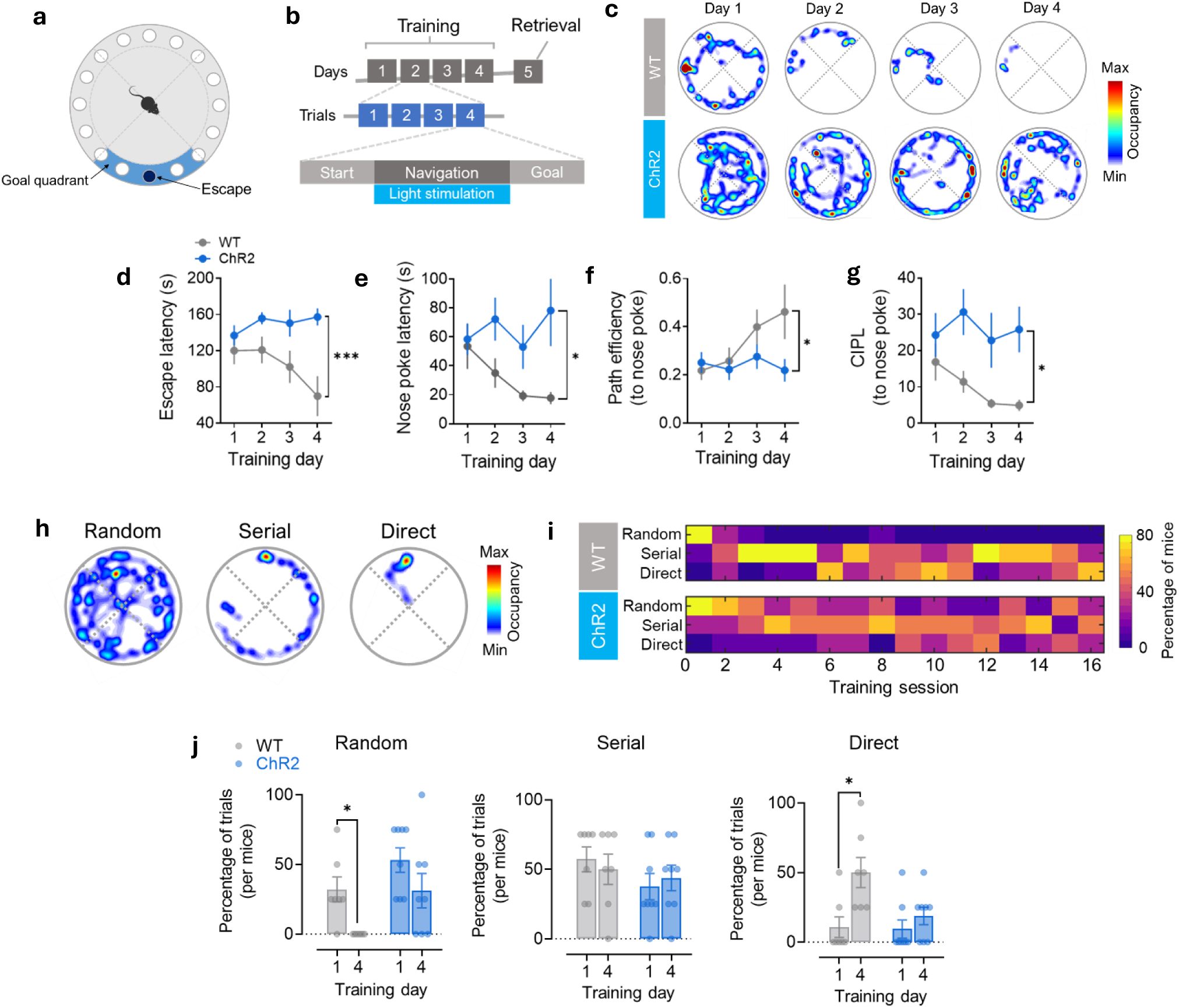
Optogenetic entrainment of neural activity in the mPFC during training in the Barnes maze impaired behavioral performance and strategy progression. **a)** Schematic diagram of the Barnes maze. **b)** Schematic diagram of the experimental design. Light stimulation was exclusively delivered during the navigation stage of all training sessions, and no light stimulation was delivered during the retrieval session. **c)** Example of color-coded occupancy plots in the Barnes maze across training days of WT (upper) and ChR2 mice (lower). **d-g)** Comparison of escape latency **(d)**, escape nose-poke latency **(e)**, path efficiency **(f),** and CIPL **(d)** between WT and ChR2 mice across training days in the Barnes maze. Data are presented as mean ± SEM. ***: P < 0.001; *: P < 0.05; Sidak’s multiple comparisons test after two-way ANOVA. **h)** Example of occupancy plots of random (left), serial (middle), and direct (right) navigation strategies in the Barnes maze. **i)** Color-surface plot comparing the percentage of WT (upper) and ChR2 (lower) mice using each of the navigation strategies across training trials. **j)** Bar and scatter plot comparing the percentage of trials in which each mouse used random (left), serial (middle), and direct (right) navigation strategies on training days 1 and 4. Each point represents a single animal. Data in the bar plot are presented as mean ± SEM. *: P < 0.05; Sidak’s multiple comparisons test after two-way ANOVA.

### 3.3. Optogenetic entrainment disrupted oscillatory dynamics and impaired the emergence of learning-associated oscillatory patterns

The spectral power of theta oscillations increased during navigation with respect to start and goal stages in the mPFC in WT mice (**Fig. 3a**). In ChR2 mice, light stimulation increased the power of theta at a similar frequency range compared to WT; this increase was significantly higher compared to WT (P = 0.032; Sidak’s multiple comparisons test after two-way ANOVA; **Fig. 3b**). The effect of optogenetic entrainment was specific for theta, as it did not affect the spectral power of 4-Hz, low and high-gamma oscillations (**suppl. Fig. 6a-c**). We then compared oscillatory dynamics. The CoV of spectral energy at the theta band was lower in ChR2 compared to WT mice, specifically during stimulation (P < 0.0001; Sidak’s multiple comparisons test after two-way ANOVA; **Fig. 3c, d**). Phase resetting, an indicative of state transition of functional networks (Canavier, 2015; Voloh & Womelsdorf, 2016), was also significantly reduced during the navigation stage in ChR2 mice compared to WT (P < 0.0001; Sidak’s multiple comparisons test after two-way ANOVA; **Fig. 3c, e**). Finally, MSE in scale factors between 30 to 120 (8 to 30 Hz) was significantly decreased in ChR2 mice compared to WT during stimulation (P < 0.01; Wilcoxon rank-sum test; **Fig. 3f, g**). We then evaluated if optogenetic entrainment affected the emergence of learning-related oscillatory patterns, such as the coupling of gamma activity by the phase of theta oscillations (i.e., cross-frequency coupling; CFC) (García et al., 2025). Optogenetic entrainment significantly affected theta-gamma coupling strength (measured as modulation index, MI) during the navigation stage (P < 0.0001; two-way ANOVA; **Fig. 3h, i**). Indeed, MI was highest in ChR2 mice compared to WT from day 1 to 3 (P < 0.05; Sidak’s multiple comparisons test; **Fig. 3i).** Interestingly, MI gradually increased in WT mice across training (P = 0.021; one-way ANOVA; **Fig. 3i; suppl. Fig. 6d**), a phenomenon not observed in ChR2 mice (P = 0.829; one-way ANOVA; **Fig. 3i; suppl. Fig. 6d**). Taken together, our analysis shows that optogenetic entrainment of prefrontal activity perturbed oscillatory dynamics exclusively during training, blocking the emergence of learning-related oscillatory patterns.

**Figure 3.**
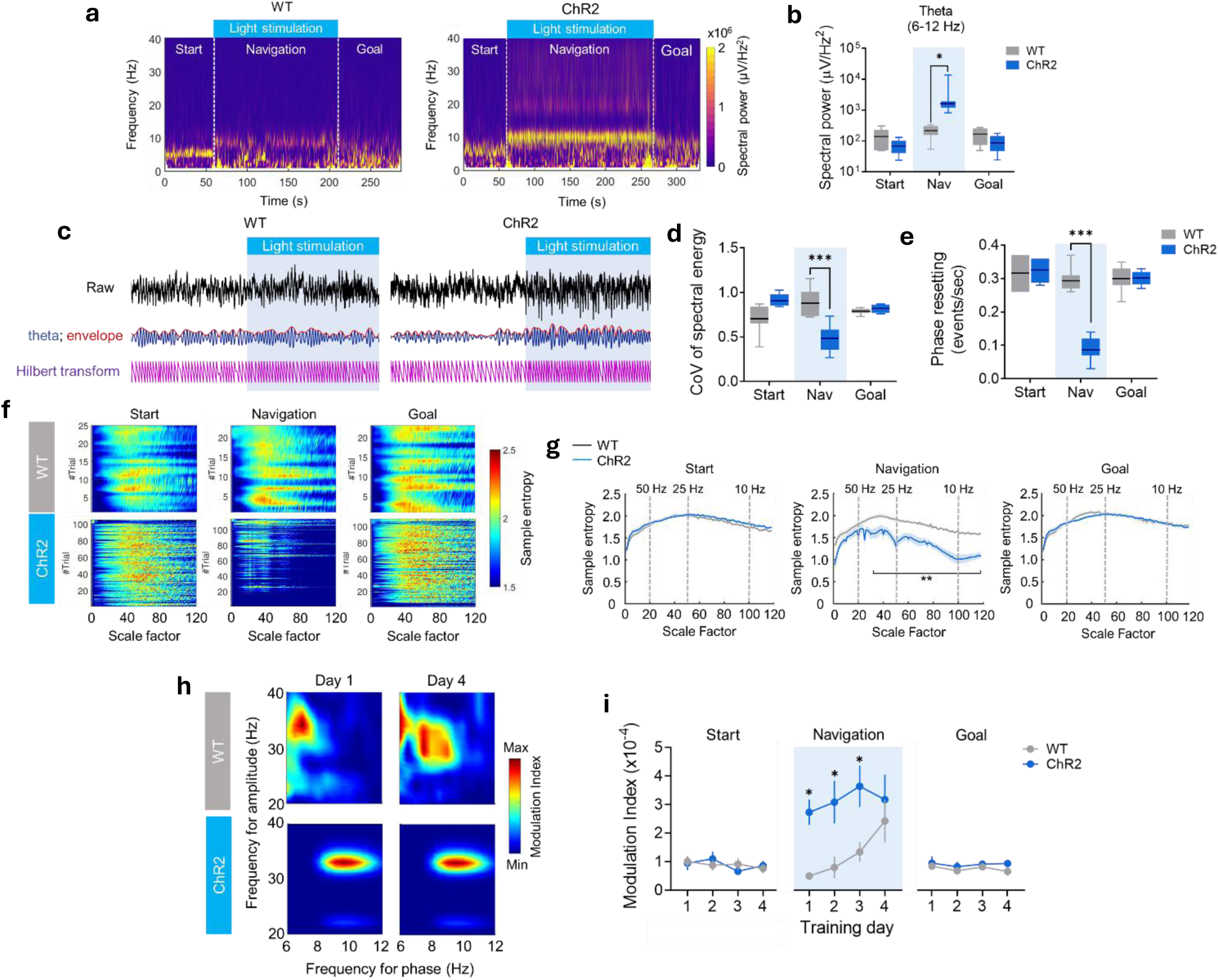
Optogenetic entrainment disrupted oscillatory dynamics and impaired the emergence of learning-associated oscillatory patterns during training in the Barnes maze. **a)** Example of time-frequency color-coded analysis of spectral power from the mPFC of WT (left) and ChR2 (right) mice during navigation in the Barnes maze. Start, navigation, and goal stages are indicated. **b)** Box plot comparing the spectral power of theta oscillations (6-12 Hz) between WT and ChR2 mice during start, navigation, and goal stages. *: P < 0.05; Sidak’s multiple comparisons test after two-way ANOVA. **c)** Example of raw (black) and theta filtered (blue) and its envelope (red) of LFP recording from the mPFC of WT (left) and ChR2 mice (right) during the transition from start to goal stages. Light illumination is indicated in both examples. **d, e)** Box plot comparing the CoV of energy of theta oscillations (**d**) and the incidence of phase resetting (**e**) between WT and ChR2 mice during start, navigation, and goal stages. ***: P < 0.001; Sidak’s multiple comparisons test after two-way ANOVA. For all the box plots, the middle, bottom, and top lines of the box plot correspond to the median, lower, and upper quartiles, and the edges of the lower and upper whiskers correspond to the 5th and 95th percentiles. **f)** Color-coded representation of MSE of all LFP signals recorded during training from the mPFC of WT (upper) and ChR2 (lower) mice during start (left), navigation (middle), and goal (right) stages. **g)** Comparison of the average MSE of the LFP signals recorded from the mPFC of WT and ChR2 mice during start (left), navigation (middle), and goal (right) stages. **: P < 0.01; Wilcoxon-signed rank test. Data are presented as mean (solid line) ± SEM (shaded area). **h)** Examples of color coded comodulograms of CFC of prefrontal low-gamma oscillations with respect to theta oscillations in WT (upper) and ChR2 (lower) mice during the navigation stage at day 1 and 4 of training. **i)** Comparison of the average modulation index (MI) of CFC from the mPFC of WT and ChR2 mice during start (left), navigation (middle), and goal (right) stages. *: P < 0.05; Sidak’s multiple comparisons test after two-way ANOVA.

### 3.4. Optogenetic entrainment disturbed the learning-related adjustment of firing patterns during training

During training, we recorded a total of 96 and 251 well-isolated single units from WT and ChR2 mice, respectively (**suppl. Fig. 7a)**. We found no differences between groups in the distribution of peak-to-valley spike waveform duration (P = 0.08; Kolmogorov-Smirnov test**; suppl. Fig. 7b, c**). We then compared firing rate between behavioral stages across groups. Given the variability of the duration of the navigation stage across trials, firing rate was calculated according to warped stage duration (Aronov et al., 2017). To this, we subdivided each behavioral stage (start, navigation, and goal) into an equal number of bins, and we calculated the normalized (z-scored) firing rate for each bin. Globally, firing increased in the mPFC of WT mice during navigation respect to start and goal stages (**Fig. 4a**) as previously reported (García et al., 2025; Negrón-Oyarzo et al., 2018). As expected, light stimulation increased firing during navigation with respect to start and goal stages in ChR2 mice (**Fig. 4a**). We found a significant effect of stages in the normalized firing (P < 0.0001; two-way ANOVA; **Fig. 4b**), which was higher in ChR2 compared to WT mice (P = 0.021; Sidak’s multiple comparisons test; **Fig. 4b**). Interestingly, also a considerable proportion of neurons decreased their firing during navigation in both WT and ChR2 groups (**suppl. Fig. 7d**). We then compared the variability of firing. Optogenetic entrainment induced rhythmic firing in ChR2 mice, as revealed by the peak at ISI of 100 ms exclusively during the navigation stage, consistent with stimulation at 10 Hz, a phenomenon not observed in WT mice (**Fig 4c**). As observed in firing rate maps (**Fig. 4d**), prefrontal firing in WT mice was progressively adjusted in the arena across training days during navigation. Contrarily, prefrontal firing in ChR2 mice remained constant during navigation, with no apparent modification across training sessions (**Fig. 4d**). To quantify these variations, we estimated the irregularity of firing for each cell by calculating the entropy of firing (Luczak, 2024; Strong et al., 1998). Entropy of firing was significantly affected by the interaction between groups and behavioral stage (P < 0.0001; two-way ANOVA; **Fig. 4e**). During navigation, the entropy of firing was higher in WT compared to ChR2 mice (P < 0.0001; Sidak’s multiple comparisons test after two-way ANOVA; **Fig. 4e**). Importantly, the entropy of firing increased across training days in WT mice (day 1 vs. day 4: P < 0.0001; Sidak’s multiple comparisons test after two-way ANOVA; **Fig. 4f**), an effect not observed in ChR2 mice (day1 vs. day 4: P = 0.997; Sidak’s multiple comparisons test after two-way ANOVA; **Fig. 4f**). On training day 1, there was no difference between WT and ChR2 mice in the entropy of firing (P > 0.999; Sidak’s multiple comparisons test after two-way ANOVA); however, at training day 4, entropy was significantly higher in WT mice compared to ChR2 mice (P < 0.0001; Sidak’s multiple comparisons test after two-way ANOVA). We then asked if the entrainment affected the emergence of task-relevant firing patterns in the mPFC. When both WT and ChR2 mice used non-spatial (i.e., random and serial) strategies, the firing of prefrontal units in turn to escape was not different from shuffling (**Fig. 4g**). Nevertheless, when WT mice used spatial strategies, we found a significant increase of firing rate briefly before entering the escape (P < 0.05; Wilcoxon rank-sum test; **Fig. 4g**), as previously documented (Negrón-Oyarzo et al., 2018). This increase of firing was not observed in ChR2 mice (P > 0.05; Wilcoxon rank-sum test; **Fig. 4g**), even when ChR2 mice scarcely implemented spatial strategies (**Fig. 2i, j**). These results suggest that optogenetic entrainment in the mPFC induced rhythmic firing, decreased firing entropy, and blocked the emergence of task-related spiking patterns.

**Figure 4.**
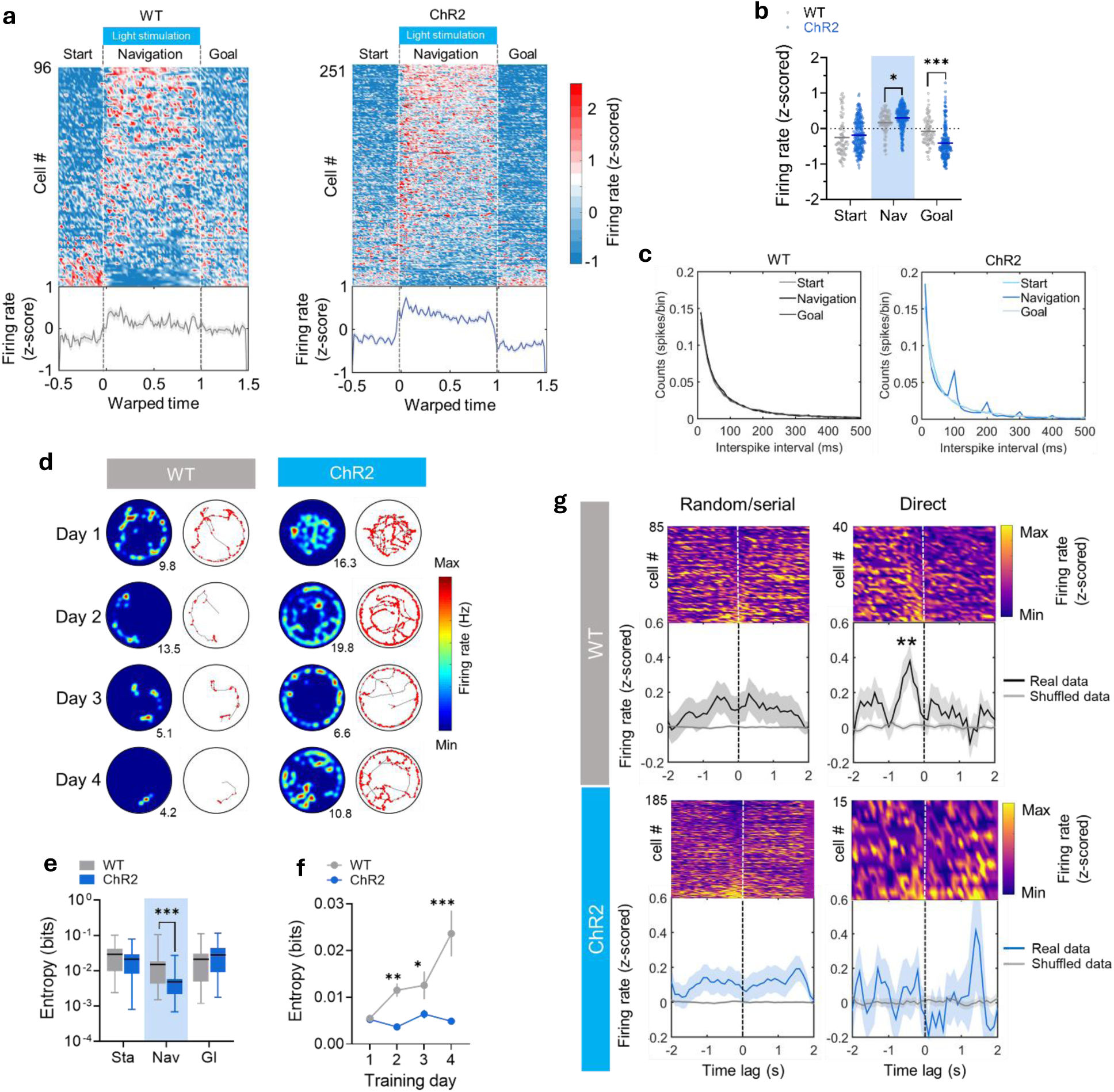
Optogenetic entrainment in the mPFC disrupted prefrontal firing dynamics and impaired the adjustment of firing patterns during training in the Barnes Maze. **a)** Upper panel: color-coded normalized firing rate (z-scored) and time (warped) for all neurons recorded in the mPFC of WT (left) and ChR2 (right) mice during all training sessions in the Barnes Maze. Start, navigation, and goal stages are indicated, as well as light stimulation. Lower panel: average peri-event time histogram of normalized (z-scored) firing rate. Shaded areas represent SEM. **b)** Scatter plot comparing the normalized (z-scored) firing rate of all units recorded from the mPFC of WT and ChR2 mice during start, navigation, and goal stages. Each point represents a single unit, and the solid line represents the mean. *: P < 0.05; ***: P < 0.001; Sidak’s multiple comparisons test after two-way ANOVA. **c**) Comparison of average inter-spike interval (ISI) between start, navigation, and goal stages in WT (left) and ChR2 (right) mice. **d)** Representative firing of prefrontal neurons across training days in the Barnes maze for WT (left) and ChR2 (right) mice. Columns on the right depict the smoothed, color-coded spatial firing rate maps scaled to the maximum firing rate (indicated in every spatial-firing rate map). Columns on the left show the path trajectories in gray and the corresponding spikes of the neurons in red. **e)** Box plot comparing the entropy of firing of neurons recorded in the mPFC of WT and ChR2 mice during start, navigation, and goal stages during training sessions in the Barnes maze. ***: P < 0.001; Sidak’s multiple comparisons test after two-way ANOVA. The middle, bottom, and top lines of the box plot correspond to the median, lower, and upper quartiles, and the edges of the lower and upper whiskers correspond to the 5th and 95th percentiles. **f)** Comparison between WT and ChR2 mice of the entropy of firing of single units recorded in the mPFC across training days in the Barnes maze during the navigation stage. ***: P < 0.001; **: P < 0.01; *: P < 0.05; Sidak’s multiple comparisons test after two-way ANOVA. Data are presented as mean ± SEM. **g)** The top of each panel shows the color-coded peri-event time histograms of the normalized (z-scored) firing rate of each cell recorded in the mPFC in turn to goal approaching, and the bottom of each panel depicts the average (the solid line represents the mean, and the shaded area represents the SEM) over all normalized firing rate. The upper panels are from neurons recorded in WT mice during random/serial strategies (upper left) and during direct navigation strategies (upper right). The upper panels are from neurons recorded in ChR2 mice during random/serial strategies (lower left) and during direct navigation strategies (lower right). **: P < 0.01*; Wilcoxon signed-rank test with respect to shuffling (gray solid line on each bottom sub-panel).

### 3.5. Optogenetic entrainment of prefrontal activity during training impaired the formation of spatial memory

It is likely that optogenetic entrainment blocked memory expression instead of memory formation (Preston & Eichenbaum, 2013). To test this possibility, WT and ChR2 mice were subjected to a single trial of spatial memory retrieval twenty-four hours after training was finalized. This trial was similar to training trials, except that light stimulation was not delivered, and the escape box was retired from the escape hole. We observed a decreased proficiency to find the escape hole in ChR2 mice compared to WT (**Fig. 5a**). Indeed, ChR2 mice showed significantly increased escape nose-poke latency (P = 0.031; unpaired t-test; **Fig. 5b**), decreased path efficiency (P = 0.023; unpaired t-test; **Fig. 5b**), a higher number of errors (P = 0.011; Unpaired t-test; **suppl. Fig. 8a**), decreased time in the goal quadrant (P = 0.037; unpaired t-test; **suppl. Fig 8b**), and increased CIPL (P = 0.035; unpaired t-test; **Fig. 8c**). Compared to WT, ChR2 mice showed a tendency to utilize more non-spatial and less spatial strategies (non-spatial: WT = 42%; ChR2 = 76%; spatial: WT = 58%; ChR2 = 24%), although these differences did not reach statistical significance (P = 0.314; Fisher’s exact test; **suppl. Fig. 8d**). These differences were not attributed to locomotor activity, as we found no differences between groups in mean and maximum speed (P = 0.613 and P = 0.316, respectively; unpaired t-test; **suppl. Fig. 8e, f**), or to genotype, as we found no differences between non-stimulated WT and ChR2 mice in escape nose-poke latency (P = 0.848; unpaired t-test), errors (P = 0.645; unpaired t-test), time in goal quadrant (P = 0.249; unpaired t-test), CIPL (P = 0.246; unpaired t-test), path efficiency (P = 0.941; unpaired t-test) or the utilization of navigation strategies (P = 0.892 Fisher’s exact test; **suppl. Fig. 8g-o**). This suggests that the entrainment of prefrontal activity during training blocked the acquisition of spatial memory. These behavioral differences could not be attributed to enduring alterations in the mPFC induced by laser stimulation during training, as we found no effect of groups in the spectral power at theta, 4-Hz, low-gamma, and high-gamma frequencies (theta: P = 0.126; 4-Hz: P = 0.284; low-gamma: P = 0.866; high-gamma: P = 0.325; two-way ANOVA; **suppl. Fig. 9a-c**). Similarly, we found no significant differences between groups in the CoV of spectral energy for theta (P = 0.084; two-way ANOVA; **suppl. Fig. 9d, e**), the incidence of phase resetting of theta oscillations (P = 0.184; two-way ANOVA; **suppl. Fig. 9d, e**) and MSE at all tested scale factors (P > 0.05; Wilcoxon rank-sum test; **suppl. Fig. 9f, g**). However, gamma-theta modulation strength was lower in ChR2 compared to WT mice during navigation (P = 0.043; Sidak’s multiple comparisons test after two-way ANOVA; **Fig. 5c, d**) effect not observed during the start stage (P = 0.878; Sidak’s multiple comparisons test after two-way ANOVA; **Fig. 5c, d**). At single unit level, we found no significant differences between groups in firing rate (P = 0.071; two-way ANOVA; **suppl. Fig. 10a, b**) and ISI (P > 0.05; Wilcoxon rank-sum test; **suppl. Fig. 10c**). However, as observed in the firing rate maps (**Fig. 5e**), neuronal firing was more regular during navigation in ChR2 mice during memory retrieval. Indeed, firing entropy was lower in ChR2 compared to WT mice during navigation (P = 0.0005; Sidak’s multiple comparisons test after two-way ANOVA; **Fig. 5f**), with no significant differences during start (P = 0.487; Sidak’s multiple comparisons test after two-way ANOVA; **Fig. 5f**). These data suggest that the entrainment during training impaired the acquisition of spatial memory and the emergence of learning-related oscillatory and firing patterns during memory retrieval, without inducing lasting effects on ongoing neural activity dynamics.

**Figure 5.**
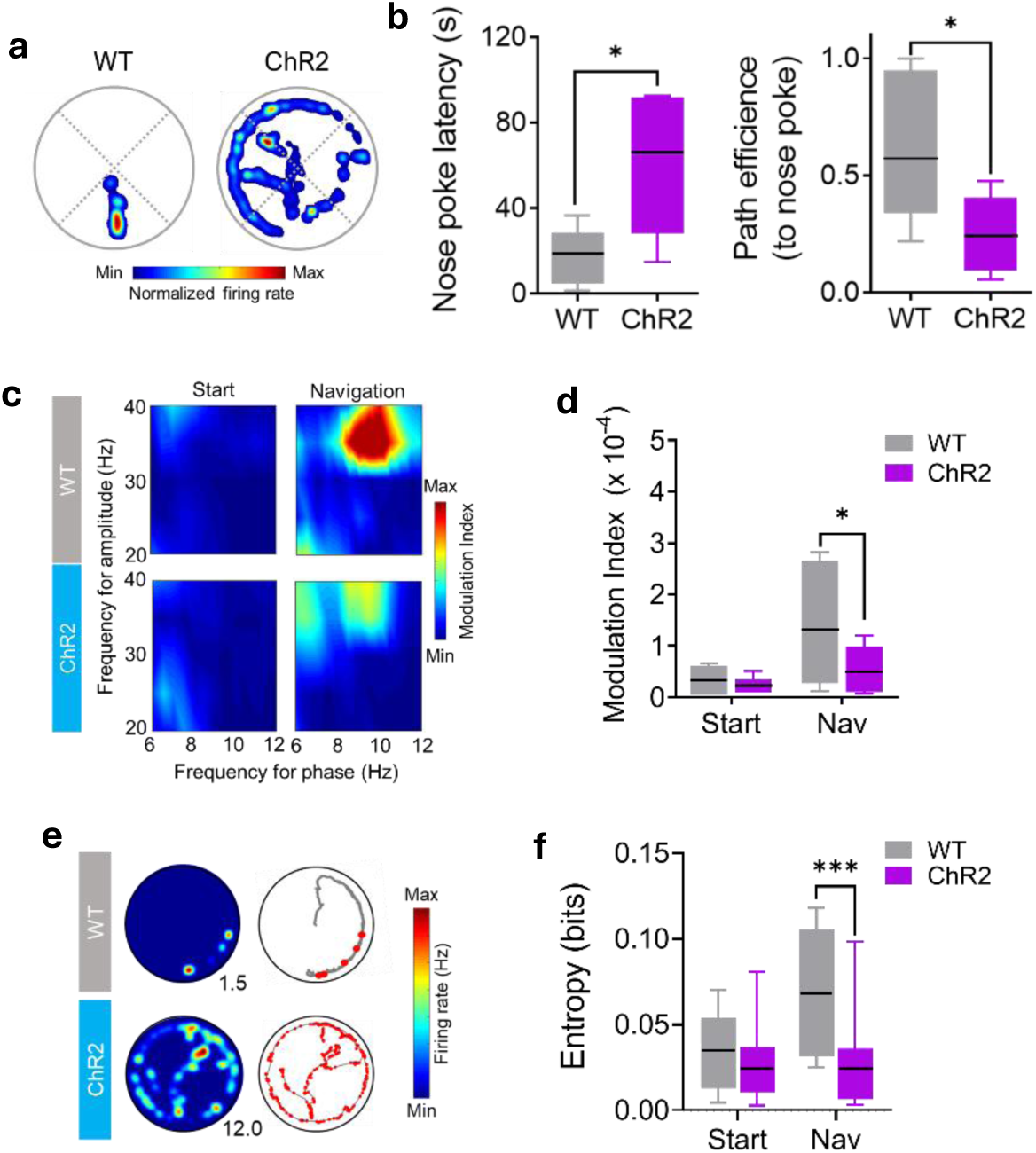
Optogenetic entrainment of prefrontal activity during training impaired behavioral performance and learning-associated prefrontal activity patterns during memory retrieval. **a)** Example of color-coded occupancy plots of WT (left) and ChR2 mice (right) during memory retrieval in the Barnes maze. **b)** Box plot comparing the escape nose-poke latency (left) and CIPL (right) between WT and ChR2 mice during memory retrieval. *: P < 0.05; Sidak’s multiple comparisons test after two-way ANOVA. **c)** Examples of color-coded comodulograms of CFC of prefrontal low-gamma oscillations with respect to theta oscillations in WT (upper) and ChR2 (lower) mice during start (left) and navigation (right) stages during memory retrieval. **d)** Box plot comparing the MI of CFC of prefrontal low-gamma oscillations with respect to theta oscillations between WT and ChR2 mice during memory retrieval. *: P < 0.05; Sidak’s multiple comparisons test after two-way ANOVA. **e)** Representative firing of prefrontal neurons during memory retrieval for WT (top) and ChR2 (bottom) mice. Columns on the right depict the smoothed, color-coded spatial firing rate maps scaled to the maximum firing rate (indicated in every spatial firing rate map). Columns on the left show the path trajectories in gray and the corresponding spikes of the neurons in red. **f)** Box plot comparing the entropy of firing of neurons recorded in the mPFC of WT and ChR2 mice during memory retrieval. ***: P < 0.001; Sidak’s multiple comparisons test after two-way ANOVA. For all box plots, the middle, bottom, and top lines of the box plot correspond to the median, lower, and upper quartiles, and the edges of the lower and upper whiskers correspond to the 5th and 95th percentiles.

## 4. DISCUSSION

Here we show that the optogenetic entrainment of neural activity at theta frequency applied in the mPFC during training reversibly disrupted local neural activity dynamics, blocked spatial learning, and hindered the emergence of activity patterns putatively required for learning. At first glance, the blockade of learning by optogenetic stimulation on excitatory neurons may seem contradictory. However, previous reports have found similar results. For example, sustained 8-Hz optogenetic stimulation of excitatory neurons in the mPFC specifically during the delay period blocked learning in the delayed non-matching-to-sample task (Liu et al., 2014). This outcome was analogous to optogenetic suppression of the mPFC, either by inhibition of excitatory neurons or by activation of inhibitory neurons (Liu et al., 2014). Similarly, low-frequency (4-5 Hz) optogenetic synchronization of excitatory neurons in macaque area V4 impaired visual discrimination (Nandy et al., 2019). This evidence raises the question about **the mechanism by which optogenetic entrainment may impair learning**. Our proposal is that optogenetic entrainment interfere with neural activity dynamics in the mPFC, blocking spatial learning. However, optogenetic stimulation may hinder learning by alternative mechanisms. For example, it has been shown that excessive optogenetic stimulation (i.e., high intensity or frequency) may suppress neuronal firing (Arenkiel et al., 2007), which may block learning (Liu et al., 2014); however, our stimulation protocol increased firing in the mPFC (**Fig. 1f**; **Fig. 4a, b**), discarding firing suppression as a putative mechanism. Also, optogenetic stimulation may prevent the shift towards the optimal oscillatory frequency required for the proper prefrontal operation (Quirk et al., 2021; Senoussi et al., 2022). Theta oscillation is the main macroscopical pattern observed in the mPFC during spatial navigation and spatial learning (Jones & Wilson, 2005; Kunz et al., 2019; Negrón-Oyarzo et al., 2018; O’Neill et al., 2013; Sirota et al., 2008). This rhythm emerges locally in the mPFC (O’Neill et al., 2013) from a mechanism likely combining hippocampal synaptic drive (Siapas et al., 2005; Sirota et al., 2008) and inhibition-induced resonance of pyramidal neurons (Stark et al., 2013). Despite the optogenetic stimulation of layer 5 pyramidal neurons at theta frequency may not be completely equivalent to naturally occurring prefrontal theta, these neurons are the main contributor to cortical LFP (Beltramo et al., 2013). Therefore, theta-frequency optogenetic entrainment by stimulation of layer 5 pyramidal neurons may resemble theta frequency coordination. As observed in **Fig. 3a**, optogenetic entrainment at theta frequency preserved theta coordination of prefrontal activity during navigation in ChR2 mice, similarly to what occurs naturally in WT mice (**Fig. 3a**); this discards the frequency shift as a putative mechanism in the blockade of learning. Finally, theta-frequency optogenetic entrainment may prevent the emergence of other oscillatory frequencies supporting learning, such as 4-Hz, and gamma oscillations (García et al., 2025). However, theta optogenetic entrainment did not affect the power of these oscillations in the mPFC during learning (**suppl. Fig. 6a-c**). Therefore, these results exclude neural activity suppression, oscillatory frequency shift, or blockade of oscillatory patterns in the hindrance of spatial learning induced by optogenetic entrainment.

Based on theoretical and analytical evidence (Deco et al., 2011; Garrett et al., 2013; Stein et al., 2005; Waschke et al., 2021), here we postulated that **optogenetic entrainment disrupt neural activity dynamics**. Our data support this statement. Whereas the energy of theta oscillations varies across navigation in WT mice, this variability was significantly reduced by optogenetic entrainment in ChR2 mice (**Fig. 3c, d**). Optogenetic entrainment also reduced the incidence of phase resetting (**Fig. 3c, e**), an estimate of state transitions in neural activity (Canavier, 2015; Voloh & Womelsdorf, 2016), and drastically reduced MSE (**Fig. 3f, g**), suggesting that optogenetic entrainment reduced the complexity of the LFP signal (Costa et al., 2005). Similar results were found in neuronal firing. Whereas firing patterns in WT animals were gradually shifting along training, neuronal firing remained uniform in optogenetically entrained ChR2 mice (**Fig 4d**). We also observed that the entropy of firing increased in WT mice as they progressed along training, a phenomenon blocked in optogenetically stimulated ChR2 mice (**Fig. 4f**). Furthermore, during memory retrieval, entropy of firing persisted high in WT mice, but not in ChR2 mice (**Fig 5f**). Importantly, our data revealed that disruption of neural dynamics was specifically restricted to the mPFC, as we found that optogenetic entrainment of prefrontal activity did not affect spectral power, CoV of theta energy, or MSE in the HPC (**suppl. Fig 4e-h**), structure connected to the mPFC (Jay & Witter, 1991; Swanson, 1981). This evidence discards the possibility that blockade of learning may be related to disruption of dynamics in other brain structures. Consequently, optogenetic entrainment of neural activity in the mPFC specifically disrupted local oscillatory and single neuron dynamics.

As a corollary of the previous statement, we also proposed that **disruption of neural dynamics might impair the development** of learning-associated activity patterns, resulting in the blockade of spatial learning. Among learning-associated oscillatory patterns, CFC seems to be particularly relevant, as it supports several neural operations, including multi-item representation and large-scale integration (Canolty & Knight, 2010; Hyafil et al., 2015). We found that local theta-gamma CFC gradually increased in WT mice along learning **(Fig. 3h, i)**, suggesting that, similarly to large-scale CFC (García et al., 2025), local CFC emerges in parallel with successful spatial learning. However, local CFC was maintained constantly high and invariant across learning in ChR2 mice (**Fig. 3h, i**). Paradoxically, CFC was pronounced in WT mice during memory retrieval, but not in ChR2 mice (**Fig. 5c, d**), suggesting that optogenetic entrainment during training prevented the formation of CFC. Similar results were found at the neuronal spiking level. We observed that firing patterns that normally emerge during spatial learning (Negrón-Oyarzo et al., 2018) emerged in WT mice; however, this was not present in ChR2 mice (**Fig. 4g**), even when spiking was enhanced by optogenetic stimulation (**Fig. 4b**). Together, these results show that optogenetic disruption of oscillatory and spiking dynamics prevented the emergence of learning-related activity patterns in the mPFC.

This evidence raises the question about **the relationship between neural dynamics and the learning-dependent development of task-associated activity patterns**. Cortical activity dynamics emerge from stochastic transitions between neural synchronization and desynchronization (Garrett et al., 2013; Harris & Thiele, 2011; Kopell et al., 2014; O’Byrne & Jerbi, 2022). These state transitions emerge from the metastability of neural activity (Hancock et al., 2024; O’Byrne & Jerbi, 2022), giving rise to the spontaneous ongoing neural activity dynamics (Buzsáki, 2006; Buzsáki et al., 2014). Thus, neural activity dynamics relies on the iteration between different configurations of reciprocal coupling between neural populations at different timescales (Buzsáki, 2006; Garrett et al., 2013; Varela, 1995). It has been proposed that the exposure of the subject to new and demanding circumstances (i.e., experience) may perturb the spontaneous neural activity, giving rise to new configurations of functional connectivity between neural populations (Buzsáki et al., 2014). This mechanism allows the task-associated integration of new information into the current cognitive operation as the subject gains experience, which is manifested as the development of particular activity patterns (Buzsáki, 2010; Buzsáki et al., 2012; Varela, 1995). In support of this, it has been observed that during learning, neurons constantly reconfigure their reciprocal coordinated activity (Agetsuma et al., 2023). Similarly, neuronal ensembles that represent relevant task features gradually develop and stabilize along learning (Baeg et al., 2003, 2007; Benchenane et al., 2010; Liu et al., 2014), which, once formed, are transiently activated according to task demands (Fujisawa et al., 2008). For this process to occur, the neural network must be configured in a labile, flexible, and receptive state that permits the reorganization of existing neural functional connectivity. Therefore, optogenetic entrainment may restrict neural networks into a fixed hyper-synchronized state, preventing the iteration among alternative arrangements of neural coordination, and thus constraining the interaction between different neuronal populations representing information potentially required for the cognitive operation. The reduction of CoV of theta energy, phase resetting, and MSE in the oscillatory activity, together with the peaks in the ISI and the decrease of entropy of spiking induced by optogenetic entrainment, agrees with the reduction of possible states of neural configurations during optogenetic entrainment. This phenomenon impacts the development of activity patterns required for task accomplishment, which agrees with the impaired development of task-related oscillatory and firing patterns along learning (**Figs. 3 and 4**). This mechanism may also explain the blockade of learning observed in other reports (Liu et al., 2014; Nandy et al., 2019).

### The present findings demonstrate the key contribution of neural activity dynamics in the mPFC in spatial learning

The mPFC is involved in the progression from less-to-more efficient navigation strategies (de Bruin et al., 1994; Kesner et al., 1989; Kolb et al., 1994; Patai & Spiers, 2021), a hallmark of spatial learning (Harrison et al., 2006; Ruediger et al., 2012). Here, we found that ChR2 mice’s strategy progression was hampered by optogenetic entrainment, which accounts for the poor performance (**Fig. 2h-j**). Strategy progression is not instantaneous, but it requires several navigation sessions over several days, in which the most efficient strategies are evident during the last training sessions (**Fig. 2i, j**). As all features of the maze remained stable during all training sessions, these behavioral changes occurred under stable environmental conditions. Therefore, it can be argued that efficient strategies are implemented once sufficient information has been collected and integrated into the cognitive operation. This implies that strategy progression depends on internal cumulative processing rather than being triggered by shifts in external demands. It is proposed that a generalized internal map-like representation is gradually constructed in the mPFC as a result of accumulated experience, which can be subsequently utilized to guide goal-directed behavior (Eichenbaum, 2017; Preston & Eichenbaum, 2013; Schlichting & Preston, 2015). Therefore, our results suggest that disruption of neural dynamics in the mPFC blocked the cognitive processing (i.e., the construction or updating of the generalized map) implied in the accumulation of information required for the proper guidance of behavior for goal achievement (Eichenbaum, 2017; Schlichting & Preston, 2015). Considering that the disruption of neural dynamics also blocked the development of learning-related patterns in the mPFC, it can be inferred that the accumulative integration of information as a product of experience is supported by the modification of the functional connectivity in the mPFC, resulting in the development of particular activity patterns supporting the cognitive operation. However, given that the mPFC is also involved in complementary immediate cognitive processes, such as attention, working memory, or memory retrieval (Euston et al., 2012; Preston & Eichenbaum, 2013), it is possible that optogenetic entrainment may affect these processes instead of the accumulation of information. We ruled out this possibility since ChR2 mice continued displaying an impaired performance during the retrieval session, similarly as during training, even in the absence of optogenetic stimulation (**Fig. 5a, b, and suppl. Fig. 8a-f**). Nevertheless, we did not discard that these “online” cognitive operations, also necessary for spatial learning, may require neural dynamics in the mPFC. In summary, here we show that neural activity dynamics is required for the development of activity patterns putatively required for proper prefrontal adjustment of behavior as a product of experience, which supports the formation of spatial memory.

Finally, it has been proposed that **neural activity dynamics** is related with normal brain function (Chialvo, 2010; Garrett et al., 2011, 2013; Grady & Garrett, 2018; Kopell et al., 2014; Stam, 2005; Voytek & Knight, 2015; Waschke et al., 2021). The proposed mechanism points out that dynamics of brain activity allow neural networks to attain a wide repertoire of alternative spatiotemporal configurations of functional connectivity. Thus, neural networks can adjust current operations and perform sudden computational transitions according to current or expected situations without reliance on modifications to the brain circuitry. Therefore, the dynamics of neural activity provide subjects continuous tuning, adaptation, and flexibility. EEG and fMRI studies in humans have shown that the dynamics of neural activity predicts memory performance and behavioral flexibility (Armbruster-Genç et al., 2016; Garrett et al., 2011), is highest in high performers and expert vs. non-expert subjects (Grundy et al., 2019) and increase with cognitive load or task complexity (Garrett et al., 2020; Grady & Garrett, 2018). Conversely, cortical dynamics is altered in several neurological and psychiatric conditions (Dinstein et al., 2015), such as autism spectrum disorder (Hecker et al., 2022), schizophrenia (You et al., 2022), and epilepsy (Protzner et al., 2010). Our findings thus contribute, with evidence from animal models, to the understanding of the role of neuronal activity dynamics in proper cognitive performance.

## 5. MATERIALS AND METHODS

### Animals

Adult male C57BL/6j (WT) and Thy1-ChR2-YFP (ChR2) mice (WT: n = 8; ChR2: n = 10; age: 60–90 days) were used in this study. Thy1-ChR2-YFP, line 18, mice were obtained from Jackson Laboratory (B6.Cg-Tg(Thy1-COP4/EYFP)18Gfng/J, stock number 007612). Mice were housed at 4 animals per cage in a 12-hour light/dark cycle in a temperature- and humidity-controlled room (22 ± 2 °C) with ad libitum access to food and water. Euthanasia was performed with an excess of inhalable isoflurane (3% mixed with O2) and confirmed by decapitation. The remains of sacrificed animals were routinely removed in special bags marked as biological waste and were compacted at the headquarters of the university before being incinerated by an external company (Veolia Chile). All experimental procedures related to animal experimentation were approved by the Institutional Animal Ethics Committee of the Universidad de Valparaíso (protocol code: BEA098-2016). Efforts were made to minimize the number of animals used and their suffering.

### Patch Clamp Experiments

Acute coronal brain slices (300 μm thick) were prepared using a vibratome (Campden Instruments, model MA752) from mice of either sex. Slices containing the mPFC were sectioned, isolated, and incubated in artificial cerebrospinal fluid (ACSF) containing (in mM): 124 NaCl, 2.69 KCl, 1.25 KH2PO4, 1.3 MgSO4, 26 NaHCO3, 10 glucose, and 2.5 CaCl2. The ACSF was adjusted to a pH of 7.4 and an osmolarity of 300–305 mmol/kg and equilibrated with 95% O2 and 5% CO2. Neurons in Layer 2/3 and Layer 5 of the mPFC were visualized using infrared differential interference contrast (IR-DIC) microscopy (Nikon, model Eclipse FN1). Whole-cell current-clamp recordings were performed with a Multiclamp 700B amplifier (Molecular Devices, Sunnyvale, CA, USA) and patch pipettes (3–6 MΩ) filled with an intracellular solution containing (in mM): 35 KMeSO4, 10 KCl, 10 HEPES-K, NaCl, 5 (Mg)-ATP, and 0.4 (Na)-GTP. The intracellular solution was adjusted to a pH of 7.2–7.3 and an osmolarity of 285 mmol/kg. For the current-voltage (I-V) protocol, a series of eight square current pulses (0.8-second duration each) were applied, starting at -100 pA with increments of +50 pA between pulses. For optogenetics in brain slices, light stimulation was controlled by a high-power LED controller, DC220 (Thorlabs, Inc.). Blue light with a nominal wavelength of 470 nm was generated using fiber-coupled LED, M470F3 (Thorlabs, Inc.). Light was delivered to the brain slices through a 200 µm fiber optic cable with a numerical aperture (NA) of 0.9, M84L01 (Thorlabs, Inc.).

### Fabrication of opto-microelectrode arrays (opto-MEA)

For simultaneous light illumination coupled with LFP and single unit recording from the mPFC, we assembled custom-made opto-MEAs carrying a single fiber optic cannula (multimode fiber, 0.39 NA, high OH, 200 μm core; catalog code: FT200UMT; Thorlabs, Inc.) surrounded by 16 tungsten wire microelectrodes (SML coated; 50 µm diameter, California Fine Wire Co., CA, USA). Opto-MEAs contained a single bundle composed of one stainless-steel tube for optic fiber (304 SS Hypo Tube, 27 ga. thin wall, .016/.0165” OD; Components Supply Co., FL, USA) surrounded by eight stainless-steel tubes for wire microelectrodes (304 SS Hypo Tube 30 ga., thin wall, .012/.0125” OD Components Supply Co., FL, USA). For the assembly of the fiber optic cannula, the fiber optics were cut to the desired length to reach the mPFC (near 1.0 mm) using a fiber optic scribe (Ruby Dual Fiber Optic Scribe; Thorlabs. Inc.) and affixed with a stainless-steel ferrule (1.25 mm multimode, LC/PC SS; Ø231 μm; Thorlabs. Inc.). For simultaneous recordings from mPFC and HPC, we incorporated an additional bundle composed of four electrode tubes to target the HPC. Laser power efficiency was measured and calibrated to every fiber optic cannula using a power light meter (PM100D Power Energy Meter; Thorlabs. Inc.). Only fiber optic cannulas with laser power efficiency higher than 80% were used in opto-MEAs. The resultant cannula was inserted into the central fiber tube of the bundle of the opto-MEA and affixed with epoxy. Microelectrode wires were cut off with iris scissors (World Precision Instruments, Sarasota Instruments), two wire microelectrodes were inserted in each electrode tube, and the length was adjusted to target the mPFC (length: 1.0 mm) and the CA1 area of the HPC (length: 1.5 mm). The final impedance of each wire microelectrode (200–500 kΩ) was measured in a saline solution at 1 kHz. Each microelectrode wire and the fiber optic canula were fixed to the tube bundle with cyanoacrylate, and wire microelectrodes were connected to a 16-channel interface board assembled with an Omnetics connector (Neurotek, Toronto, ON, Canada).

### Opto-MEA implantation surgery

Animals of both groups were anesthetized with isoflurane (3% isoflurane with 0.8% O2) before being placed in a stereotaxic frame. Anesthesia was maintained until the surgery was finished (1–2% isoflurane with 0.8% O2). After an incision in the scalp, one craniotomy was drilled in the right hemisphere at stereotaxic coordinates targeting the mPFC (1.94 mm AP, -0.25 mm ML, from Bregma; (George Paxinos, 2001). For the implantation of the opto-MEA targeting simultaneously the mPFC and HPC, an additional craniotomy was drilled to reach the HPC (coordinated from Bregma: -1.94 mm AP, -1.5 mm ML; (George Paxinos, 2001)). The dura was removed, and the opto-MEA was inserted into the craniotomy, reaching the cortical surface, and was slowly lowered (0.2 mm/min) until the electrodes and the fiber optic were completely inserted into the desired area. Two ground wires were attached to skull screws. Once in position, the opto-MEA and the ground screws were affixed to the skull with dental acrylic. After surgery, animals were maintained in individual cages in a temperature- and humidity-controlled room (22 ± 2 °C) with food and water *ad libitum* and were supplied with a subcutaneous dose of analgesics (Ketoprofen, 5 mg/kg/day) and antibiotics (Enrofloxacine, 5 mg/kg/day) during the 5 days after surgery. Mice were allowed to recover for at least one week after surgery before the beginning of testing and behavioral experiments. During recovery, weight and general health were monitored daily.

### Light power intensity testing

After recovery from surgery, we tested the minimal light intensity to evoke firing in the mPFC of each chronically implanted ChR2 mouse. To this aim, every individual opto-MEA implanted mouse was placed in an open field (40 cm x 40 cm x 40 cm); the fiber optic cannula of the opto-MEA was connected via a fiber optic rotary joint (Pigtail Rotary Joint, 200um 0.39NA Fiber, FCPC, 1.25, Ferrule; Thorlabs. Inc.) to the laser delivery system (LRS-0473-PFM-00050-05, Lab Spec, 473nm, DPSS Laser System; Laserglow Technologies, ON, Canada), and the interface board of the opto-MEA was connected to the amplifier board (RHD2000 evaluation system; Intan Tech, CA, USA) via a 16-channel head-stage (model RHD2132; Intan Tech, CA, USA). The laser delivery system was controlled via TTL either by a USB pulse train generator (Pulser, PRIZMATIX LTD) or an arbitrary function generator (AFG-2000 Series; GW Instek Instrument Co., Ltd). The optical stimulation system was coupled and synchronized to the electrophysiological recording system. Once the mouse was connected, they were allowed to freely explore in the open field for 1 minute with the laser off (pre stage). Then, the laser was turned on at the desired power intensity, and mice were left to explore for 1 min (stimulation stage); then the laser was turned off, and the mouse was left to explore in the maze for another minute (post stage). This procedure was repeated for each laser intensity tested. Laser stimulation consisted of pulses of 15 ms duration, separated by 85 ms between pulses, granting a sustained 10 Hz stimulation at 0.0, 0.1, 0.5, 1.0, 1.5, and 2.0 mW power laser intensities. The same procedure was achieved with WT mice. Neural activity was recorded during the entire procedure, digitized at 20 kHz, filtered at 0.1-5 kHz, monitored through the interface software (Intan Tech, CA), and stored in a hard disk for offline analysis.

### Memory acquisition task and electrophysiological recording procedures

The next day after testing, mice were individually habituated and placed in the behavioral room for 15 minutes, and then they were trained for the spatial reference memory task in the Barnes maze (Barnes, 1979; Harrison et al., 2006). The maze consisted of a white circular platform of 70 cm in diameter elevated at 70 cm from the floor with 16 equally spaced holes (9 cm in diameter) along the perimeter and located at 2 cm from the edge of the platform. Visual cues were located on the walls of the room. Under one of the holes was located a black plexiglass escape box (17 x 13 x 7 cm) that allowed the mice to enter with the implanted opto-MEA. The spatial location of the escape box was consistent for a given mouse but randomized across the mice group. The maze was illuminated with two incandescent lights to yield a light level of approximately 400 lux impinging on the circular platform. To evaluate the acquisition of spatial memory (training), mice were subjected to four consecutive behavioral sessions per day with an inter-trial interval of 15 minutes during four consecutive days. In each training session, the fiber optic cannula and the interface board of the opto-MEA were connected to the laser delivery system and the amplifier, respectively, similarly to the laser intensity testing. Then the recording system was turned on, and the mouse was placed in the start box in the center of the maze for 1 min with the room lights turned off (start stage). After time had elapsed, the start box was removed, the room lights turned on, the laser delivery system was turned on (1.0 mW of power intensity), and the mouse was free to explore the maze (navigation stage). The navigation stage ended when the mouse entered the escape box or after 3 minutes elapsed. Once the mouse entered the escape, the lights were turned off, as well as the laser delivery system, and the mouse was left to remain in the escape box for 1 min (goal stage). If the mouse did not enter the escape hole within 3 minutes, the experimenter guided the mouse to the escape. When the session was completed, the electrophysiological recording system was turned off, the mouse was disconnected from the electrophysiological recording and light stimulation system, and returned to the home cage. Then, the maze was cleaned with 70% ethanol to prevent a bias based on olfactory or proximal cues within the maze. Twenty-four hours after the training was completed, a memory retrieval session was conducted. This was similar to training trials, except that the escape box was retired from the maze and that the laser system was off. The animal behavior was tracked and recorded during each training and retrieval session with a webcam (acquisition at 30 fps and 640 × 480 pixels; camera model C920; Logitech Co.) located 1 m above the maze and controlled with VirtualDub software. Electrophysiological recordings and behavioral videos were stored on a hard disk for offline analysis. For all behavioral sessions, the electrophysiological recording system was coupled and synchronized with the optical stimulation and behavioral recording system. This same behavioral procedure was applied for the evaluation of spatial memory acquisition in mice non-implanted with the opto-MEA (n = 9 per group).

### Histology

After the experiment was finished, each mouse was anesthetized with 3% isoflurane and then transcardially perfused with ∼40 ml of phosphate-buffered saline (PBS, pH = 7.4), followed by ∼50 ml of paraformaldehyde (PFA) in PBS. The brain was removed and then stored in PFA with PBS overnight. PFA was washed with a solution of PBS containing 30% saccharose, and brains were stored in PBS with 30% saccharose until the cut session. Coronal brain slices (50 µm) from the mPFC were obtained from a cryostat-microtome (Zhejiang Jinhua Kedi Instrumental Equipment Co., LTD) and stored in PBS containing 0.01% sodium azide. Brain sections were processed with DAPI staining and observed under fluorescence microscopy (Ex: 488 nm; Nikon). Phenotyping was confirmed by the presence of yellow-fluorescent protein (YFP) on cortical layer 5 pyramidal neurons.

### Behavioral analysis

#### Behavioral performance

Behavioral performance was analyzed by measuring the escape latency (the time to enter the escape hole), escape nose-poke latency (time elapsed for the first contact of the mouse nose with the escape hole), number of errors (nose-pokes in non-escape holes), distance covered, path efficiency, and corrected integrated path length (CIPL) to the first nose-poke in the escape hole. To assess locomotor activity, the mean and maximum speed were measured in all trials. All these behavioral parameters were analyzed and measured with ANY-maze software (Stoelting Co.). Also, ANY-maze software provided for every trial a file containing instantaneous spatial coordinates of the animal’s trajectory and instantaneous speed. These were used to construct occupancy plots and spatial firing rate maps using MATLAB software with custom-made scripts.

#### Identification of navigation strategies

Navigation strategies were defined and analyzed with the Barnes Maze Unbiased Strategy Classification Tool (BUNS) (Illouz et al., 2016) implemented in MATLAB software. Coordinates of the animal trajectory obtained from ANY-maze software were entered in MATLAB by using the support vectorial machine (SVM) and converted into Cartesian coordinates to characterize animal trajectory features until the first contact with the escape hole. Data were adjusted at BUNS detector requirements, and then the strategies were classified according to the following definitions: random, when the animal navigates randomly in the maze until it finds the escape hole; serial, when the animal navigates from hole to hole (adjacent holes) until it finds the escape hole; focused search, when the animal realizes a located scan of the escape hole quadrant; long correction, when the animal navigates to an erroneous hole, and then it adjusts the navigation trajectory to directly find the escape hole; corrected, when the animal slightly modifies its trajectory (go to the next hole, before the escape hole); direct, when the animal uses the shortest path to find the escape hole. All the results obtained from the strategy detector were compared with a visual analysis recording to determine possible errors.

#### Electrophysiological analysis

##### Power spectral density analysis of the LFP

For analysis of the spectral power of oscillatory activity, electrophysiological recordings were down-sampled to 1000 Hz and bandpass-filtered at 0.1–120 Hz. Power spectral density (PSD) was computed using multi-taper Hamming analysis using the Chronux toolbox (http://www.chronux.org; (Mitra & Pesaran, 1999)), for which LFP recordings were divided into 4000 ms segments with 400 ms overlap and a time-bandwidth product (TW) of 5-9 tapers. This analysis was performed with custom-made scripts written in MATLAB software.

##### Coefficient of Variation of Instantaneous Energy

For the estimation of variability of spectral energy of theta oscillations, the down-sampled LFP recordings were filtered at theta frequency (6-12 Hz) using the Hilbert transform, and the resultant envelope (i.e., instant energy) was rectified, smoothed, and z-scored (Belitski et al., 2010). We then calculated the coefficient of variability of the resultant instant energy for every stage according to the TTL signal indicating the LASER on and off. This analysis was performed with custom-made scripts written in MATLAB software.

##### Phase resetting events detection

A finite impulse response (FIR) filter in the theta band (6-12 Hz) was applied to the raw LFP (downsampled to 1000 Hz) and the Hilbert transform instantaneous phase was calculated. Individual theta cycles were defined as intervals between phase transitions crossing -6 radians. Putative phase resetting events were identified as instances where intracycle phase differences negatively exceeded a threshold of -0.1 radians.

##### Multiscale entropy

Multiscale entropy (MSE) calculates the sample entropy (SE) of a signal at different coarse-grained temporal scales (Costa et al., 2005). To this aim, the signal is first coarse-grained into different pre-established scales, and then the SE was calculated for each of these scales. For the first step, consecutive coarse-grained time series (y^τ^) are constructed by averaging the data points within non-overlapping windows of length τ. Thus, the length (number of points) of each coarse-grained time series is equal to the length of the original time series divided by the scale factor, τ. This is expressed in the following equation:

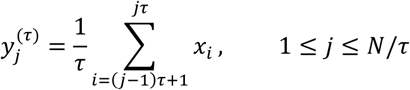

Therefore, from the acquisition rate of the time data series, each coarse-grained data series corresponds to the signal at a given frequency (Fq) as described by the following equation:

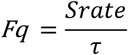

Here we established the Srate = 1000 Hz, and τ = 120. This allows us to calculate MSE to frequencies up to 8 Hz. Then the SE, which estimates the degree of randomness (or inversely, the degree of orderliness), was calculated for each coarse-grained time series. Each time data series is broken into vectors of length “m” and moved forward “τ” time points each step. Each resultant vector is compared to all other vectors except themselves, and similarity between vectors (p) is established according to the tolerance “r”. Thus, this procedure counts how often patterns of successive data points reoccur in time. This same procedure is also calculated using vectors of length “m+1”. SE is the ratio between matches on a vector of length “m” with respect to matches on a vector of length “m+1”, as described in the following equation:

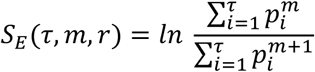

Consequently, SE estimates the temporal regularity as many patterns of length m are not repeated at length m+1. Here we established r = 0.15, m = 2, and m = 3000, as recommended (Costa et al., 2005). MSE of the LFP was quantified using a custom-made script written in MATLAB software based on the function “multiscaleSampleEntropy_compatible” (John Malik, 2025. Multiscale Sample Entropy, MATLAB Central File) based on (Costa et al., 2005)).

##### Cross-frequency coupling analysis

For CFC analyses, the ModIndex toolbox (Tort et al., 2010) was used with custom-made scripts in MATLAB software. CFC was computed using the Modulation Index (MI). The MI is able to detect the strength of the phase-amplitude coupling between two frequency ranges of interest: the “phase-modulating” (theta) and “amplitude-modulated” (gamma) frequency bands. First, LFP was filtered at the two frequency ranges under analysis. Next, the phase and the amplitude time series were calculated from the filtered signals by using the Hilbert transform. Specifically, the time series of the phases were obtained from the standard Hilbert transform, and to extract the amplitude envelope, we also applied the Hilbert transform. Then, the composite time series was constructed, which gave us the amplitude of the “amplitude-modulated” oscillation at each phase of the “phase-modulating” oscillation. The phases were binned, and the mean amplitude over each phase bin was calculated. The existence of phase-amplitude coupling is characterized by a deviation of the amplitude distribution P from the uniform distribution U in a phase-amplitude plot (if there is no phase-amplitude coupling, the amplitude distribution P over the phase bins is uniform). A measure that quantifies the deviation of P from the uniform distribution was determined by an adaptation of the Kullback-Leibler (KL) distance (DKL), which is widely used in statistics to infer the amount of difference between two distributions. The adaptation was to make the distribution distance measure assume values between 0 and 1. The MI is therefore a constant time, the KL distance of P from the uniform distribution (U). Thus, the higher the MI, the greater the phase-amplitude coupling between the frequencies and areas of interest. It is represented in the following equation:

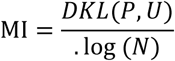

##### Spike sorting

Spike sorting was performed offline using Wave-clus, a MATLAB-based super paramagnetic clustering method (Quiroga et al., 2004). To detect spikes, broadband recordings sampled at 20 kHz were filtered at 500–5000 Hz, and events with amplitudes reaching between 4 and 8 standard deviations over the mean were considered candidate spikes. The spike features considered for clustering included peak amplitude (the maximum height of the waveform of each channel of each spike), timestamp (the time of occurrence of each spike), spike width (the duration in each channel of the spike), energy (the energy contained within the waveform of each channel of the spike), valley (the maximum depth of the waveform of each channel of the spike), and wave-PC (the contribution to the waveform due to the principal component). A file for every single unit containing the timestamp of all detected spikes was generated and used in the subsequent analyses on MATLAB. To confirm the correct sorting of single units, the time stamps of every single unit were aligned with their 500–5000 Hz filtered LFP recording, and the concordance of timestamps with spikes was visually established. To distinguish the neural identity of single units (putative regular from fast spiking units), the timestamp of every single unit was aligned with its 500 Hz high-pass filtered LFP recording, and maximum and minimum peaks were detected for every spike. Next, we obtained an average spike waveform that considered all the spikes of each unit, and we calculated the valley-to-peak (VTP) duration. Units with VTP duration lower than 0.4 ms were considered narrow spike units (putative fast spiking), and those higher than 0.4 were considered wide spike units (putative regular spiking).

##### Analysis of spiking rate between stages

The firing rate of each single unit on different behavioral stages was calculated by dividing the number of timestamps of spikes by the duration of the stage. Given the differences in firing rate between single units, the comparison of firing rate between stages was realized as a normalized z-score. For the calculation of the inter-spike interval of each unit across stages, we calculated the time differences between adjacent spikes, and we obtained the distribution (bin = 10 ms) of these values for each unit for different stages. For warping analysis, each behavioral stage was subdivided into an equal number of bins, and the z-scored firing rate was calculated for each bin.

##### Spatial-firing rate maps

Coordinates of animal trajectory obtained from ANY-maze software were loaded in MATLAB and converted into Cartesian coordinates. The space was divided into 740 x 570 pixel bins (1 bin = 1.58 cm). The timestamp of spikes was converted into a three-dimensional histogram of bivariate data in MATLAB, and spikes were signaled in the maze space. The firing rate for each bin was calculated and smoothed using two-dimensional convolution followed by the Hanning method.

##### Entropy of spike trains

For the estimation of the entropy of firing, we first calculated the time differences between adjacent spikes for every single unit timestamp, and the Shannon entropy (Shannon, 1948) was calculated for the distribution of time differences between spikes as described in the following equation:

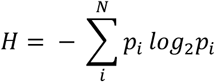

Where H is the entropy, N is the number of total events (time differences between spikes), and pi is the probability of the *i*th event.

##### Peri-event time histograms

Activity of prefrontal neurons in turn to events of behavioral relevance was evaluated by cross-correlation applying the “sliding-sweeps” algorithm (Abeles & Gerstein, 1988). First, the timing of behaviorally relevant events, such as escape (entering the escape hole) was established by analyzing the recorded videos of behavioral performance using ANY-Maze software. Then, a time window of ±2 s was defined with the 0-point assigned to the start time of the assigned behavioral event. The timestamps of the prefrontal cortex neuronal spikes within the time window were considered as a template and represented by a vector of spike numbers relative to t = 0 s, with a time bin of 100 ms and normalized to the total number of spikes. Thus, the central bin of the vector contained the ratio between the number of prefrontal cortex neuronal spikes elicited between ± 100 ms and the total number of spikes within the template. Next, the window was shifted to successive assigned behavioral events throughout the recording session, and an array of recurrences of templates was obtained. Both prefrontal cortex neuronal spike timestamps and start times of behavioral events were shuffled by randomized exchange of the original inter-event intervals (Nádasdy et al., 1999) and the cross-correlation procedure was performed on the pseudo-random sequence. The statistical significance of the observed repetition of spike sequences was assessed by comparing, bin to bin, the original sequence with the shuffled sequence. An original correlation sequence that presented a statistical distribution different from 100 simulated shuffling was considered statistically significant, with a P < 0.01 probability, instead of a chance occurrence.

### Statistical analysis

Comparisons between the means of quantitatively normally distributed variables in two groups (such as behavioral parameters during memory retrieval) were analyzed using the t-student test. Comparisons between the means of quantitatively non-normally distributed variables in two groups (such as basal firing rate between groups) were analyzed using the Mann-Whitney test. Comparisons between the means of quantitatively normally distributed variables of more than two groups (such as behavioral parameters across strategies) were analyzed using one-way ANOVA followed by Tukey’s multiple comparisons. Comparisons between the means of quantitative variables across two categorical independent variables (such as most behavioral and neurophysiological parameters) were performed using a two-way ANOVA followed by Sidak’s multiple comparisons test. Distributions (as nosepoke duration values across training days) were compared using the Kolmogorov-Smirnov test. Comparisons between the means of two quantitatively normally distributed matched samples (such as MSE across scale factors) were performed using the Wilcoxon signed-rank test. Statistical analysis was performed with GraphPad Prism or MATLAB (The Mathworks Inc.) software. Significant differences were accepted at P < 0.05.

## 7. ACKNOLEDMENTS

This work was supported by ANID Chile, projects FONDECYT INICIACION N°11160251 and ANILLO ACT210053 to I.N-O., and Beca de DOCTORADO to T.D and L.C-V. I.N-O acknowledges the support of the Universidad de Valparaíso: INICI-UV (UVA20993), and Centro de Neurobiología y Fisiopatología Integrativa (DIUV-CI 01/2006). MF acknowledges the support of FONDECYT REGULAR N°1241173 and CIDI n°1 CENFI-UV. We thank Dra. Francisca Garcia for her valuable comments in the first version of the manuscript. We thank Dr. Pablo Moya, Dra. Ana María Cárdenas, Dr. Alexies Dagnino, and Dr. Ramón Sotomayor for sharing physical space and equipment for histological experiments.

## 8. AUTHOR CONTRIBUTIONS

I.N-O. conceived the idea, designed the experiments, provided funding, interpreted the data, and wrote the manuscript. I.N-O., T.D., L.C-V., and D.B. carried out the *in vivo* experiments. I.N-O. and N.E. developed analytical scripts. D.B. analyzed behavioral data. I.N-O. analyzed electrophysiological data. D.B. analyzed histological data. K.M and M.F carried out the *ex-vivo* slice experiments. All authors discussed and commented on the manuscript.

## SUPPLEMENTARY FIGURES

**Supplementary figure 1.**
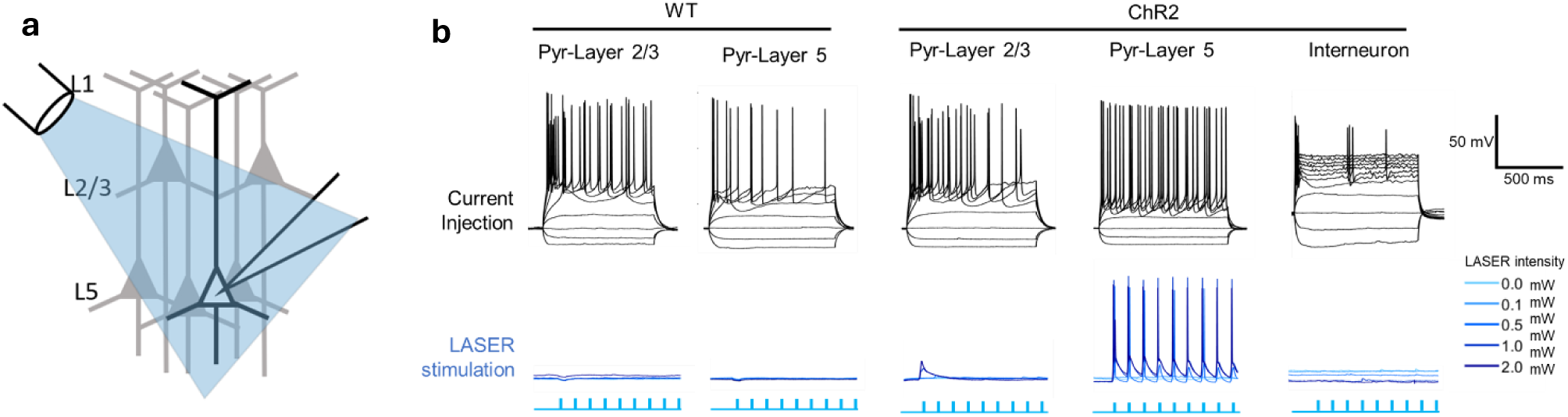
**a)** Schematic diagram showing the combined patch clamp recording with light stimulation. **b)** Representative voltage traces of patch clamp recordings (configuration current clamp) from pyramidal neurons from layers 2/3 and 5, and an interneuron from slices containing the mPFC from WT and ChR2 mice. Upper panel: voltage responses to hyper- and depolarizing current pulses. Lower panel: voltage responses to 10 Hz light stimulation at different light power intensities.

**Supplementary figure 2.**
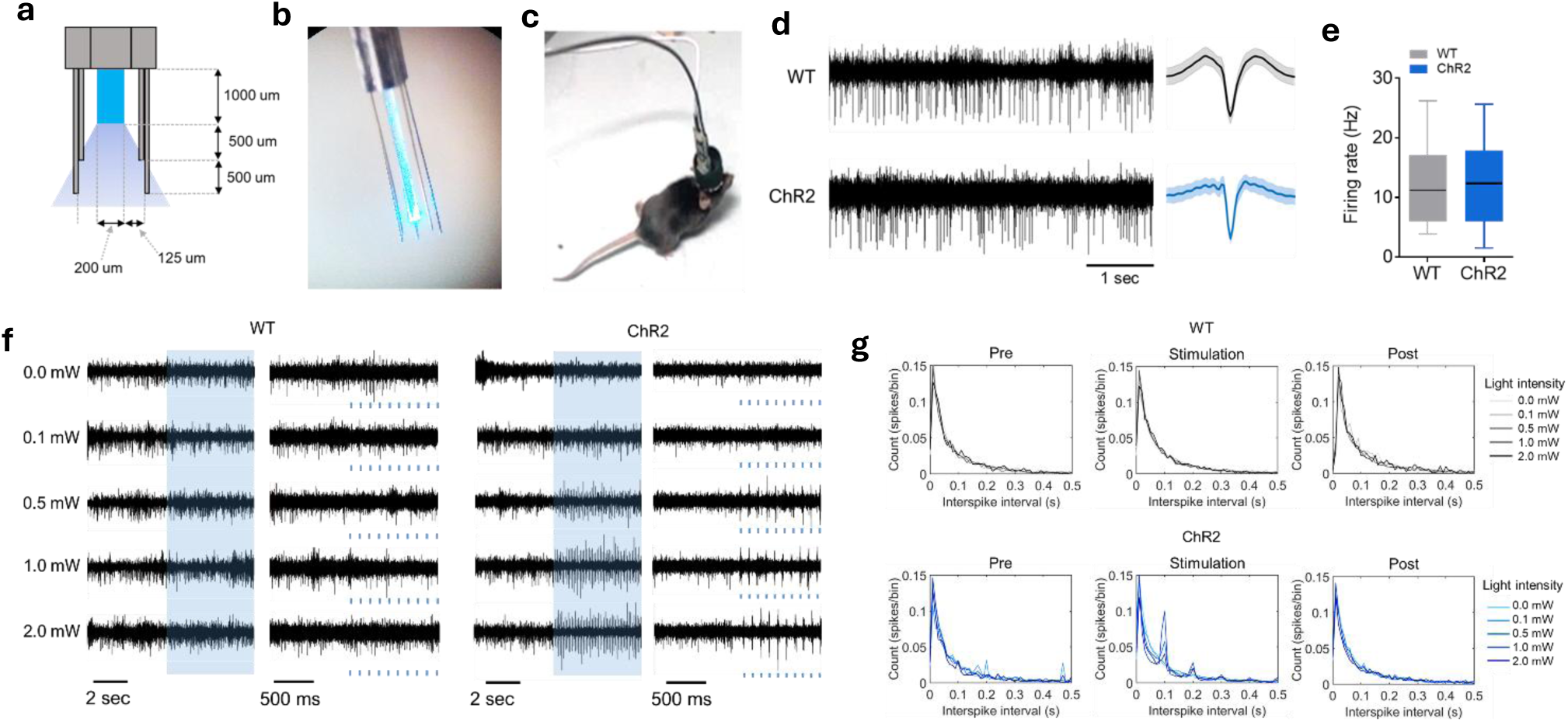
**a, b)** Schematic diagram (**a**) and photography (**b**) of the arrangement of the fiber optic and wire microelectrode in an opto-MEA. **c)** Photography of mouse chronically implanted opto-MEA and connected to the electrophysiological recording and light-delivery system. **d)** Representative recording of a single unit (LFP filtered at 500-5000 Hz) from the mPFC of a WT and ChR2 mouse. The average waveform for each single unit is shown on the right. **e)** Box plot comparing the basal firing rate (without light stimulation) of all single units detected in the mPFC of WT and ChR2 mice during testing. The middle, bottom, and top lines of the box plot correspond to the median, lower, and upper quartiles, and the edges of the lower and upper whiskers correspond to the 5th and 95th percentiles. **f)** Representative single unit recording from the mPFC of WT and ChR2 mice during the transition from laser-off to laser-ON at different light intensities during the testing sessions. The column on the right is a magnification of the column on the left. **g)** Comparison of the average inter-spike interval (ISI) between single units recorded from the mPFC of WT and ChR2 mice between pre, stimulation, and post stages at different light power intensities used during the stimulation stage of the testing sessions.

**Supplementary figure 3.**
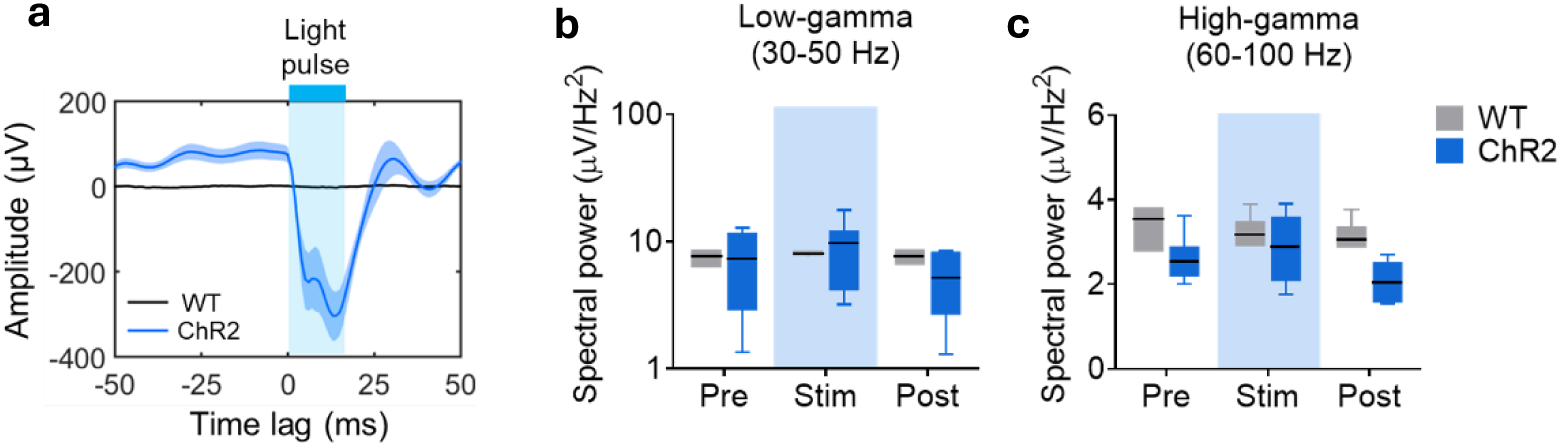
**a)** Comparison of the amplitude of light-evoked potential recorded from the mPFC of WT and ChR2 mice at power light intensity of 1 mW. Data are presented as mean (solid line) ± SEM (shaded area). **b-c)** Box plot comparing spectral power of low-gamma (**b**) and high-gamma (**c**) oscillations in the mPFC of WT and ChR2 mice across pre, stimulation, and post stages of the testing sessions. The middle, bottom, and top lines of the box plot correspond to the median, lower, and upper quartiles, and the edges of the lower and upper whiskers correspond to the 5th and 95th percentiles.

**Supplementary figure 4.**
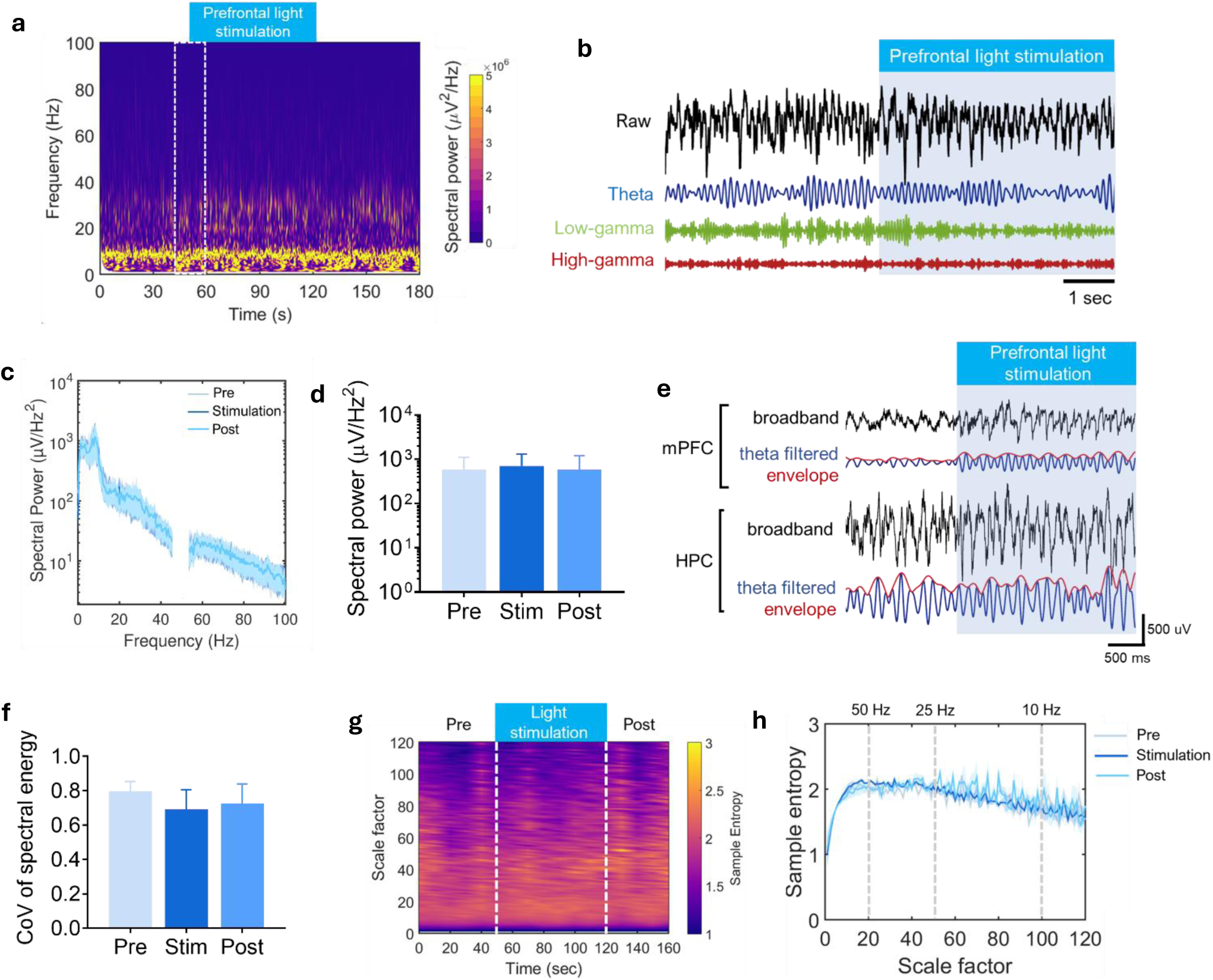
**a, b)** Example of time-frequency color-coded analysis of spectral power (**a**) and broadband and filtered (4-Hz, theta, low-gamma, and high-gamma) LFP recording (**b**) from the HPC of a ChR2 mouse during 10-Hz light stimulation in the mPFC in the transition between pre-off and stimulation-on during a testing session. Light stimulation in the mPFC is indicated in both examples. **c, d)** Average power spectral density analysis **(c)** and bar chart comparing the spectral power of theta oscillations (**d**) of the LFPs recorded from the HPC of ChR2 mice during pre, stimulation and post stages of the testing sessions. Data in **(c)** are presented as mean (solid line) ± SEM (shaded area); data in (**d**) are presented as mean ± SEM. **e)** Example of broadband (black) and theta filtered (blue) and its envelope (red) of simultaneous LFP recordings from the mPFC (upper) and HPC (lower) of a ChR2 mouse during the transition from pre-off to stimulation-ON stages during a testing session. Light stimulation in the mPFC is indicated. **f)** Bar chart comparing the CoV of spectral energy of theta oscillations during pre, stimulation, and post stages of the testing sessions. Data are presented as mean ± SEM. **g)** Example of time-scale analysis of multiscale entropy (MSE) of LFP signal recorded from the HPC of ChR2 mice during 10-Hz light stimulation in the mPFC. **h)** Comparison of the average MSE of the LFP signals recorded from the HPC of ChR2 mice across pre, stimulation and post stages of the testing sessions. Data are presented as mean (solid line) ± SEM (shaded area).

**Supplementary figure 5.**
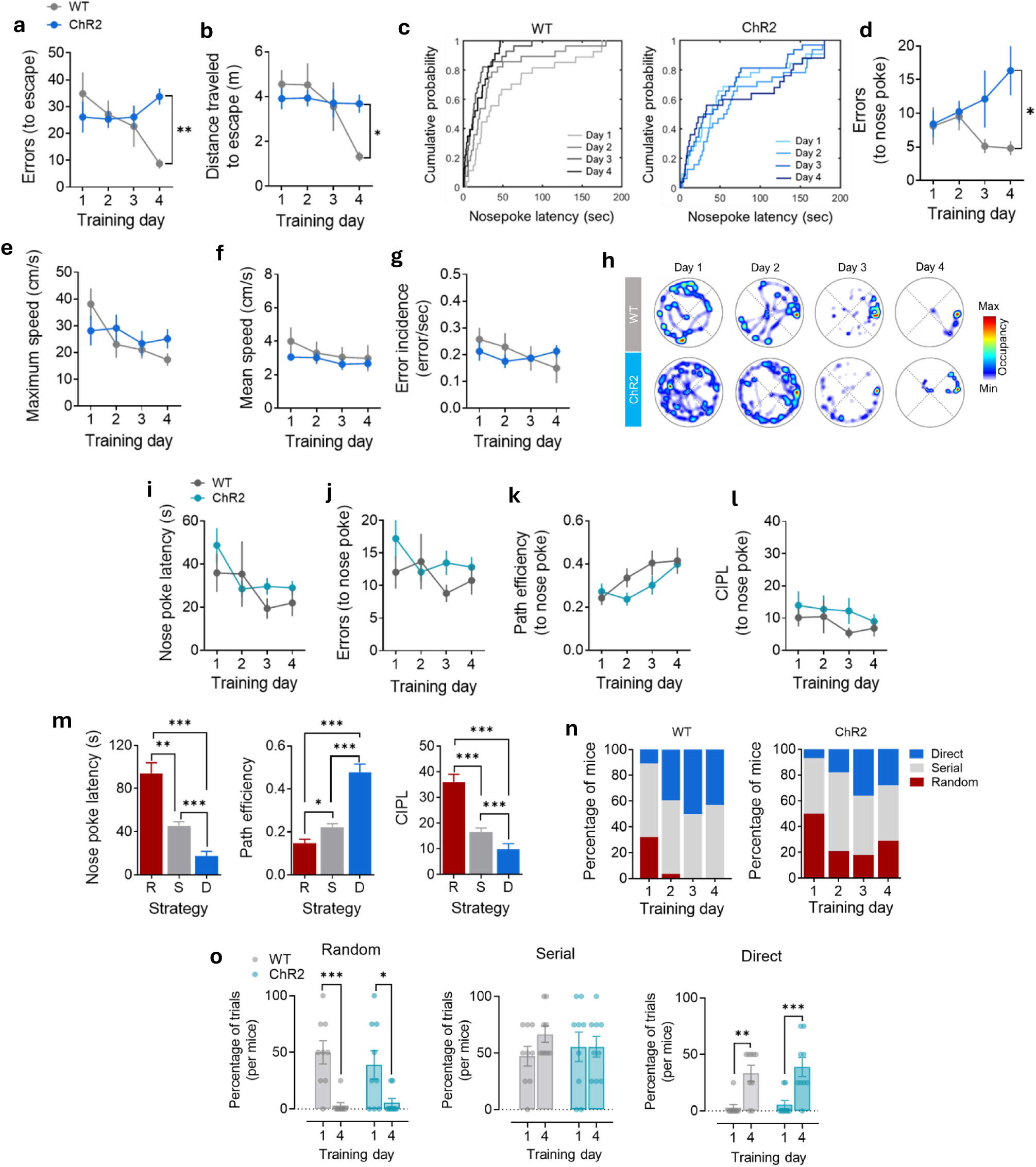
**a, b)** Comparison of the average errors (**a**) and distance (**b**) to escape between WT and ChR2 mice across training days in the Barnes maze. Data are presented as mean ± SEM. **: P < 0.01; *: P < 0.05; Sidak’s multiple comparisons test after two-way ANOVA. **c)** Cumulative histogram comparing the distribution of the duration of nose-poke latencies across training days of WT (left) and ChR2 (right) mice. **d-g)** Comparison of the average errors to nose-poke (**d**), maximum speed (**e**), mean speed (**f**), and error incidence (**g**) between WT and ChR2 mice across training days in the Barnes maze. *: P < 0.05; Sidak’s multiple comparisons test after two-way ANOVA. Data are presented as mean ± SEM. **h)** Example of color-coded occupancy plots in the Barnes maze across training days of non-light stimulated WT (upper) and ChR2 mice (lower). **i-l)** Comparison of the average latency (**i**) errors (**j**) path efficiency (**k**) and CIPL (**l**) to escape nose-poke between non-light-stimulated WT and ChR2 mice across training days in the Barnes maze. Data are presented as mean ± SEM. **m)** Bar chart comparing average escape nose-poke latency (left), path efficiency (middle), and CIPL (right), across navigation strategies during training in the Barnes maze (R: random, S: serial, D: direct). ***: P < 0.001; **: P < 0.01; *: P < 0.05; Tukey’s multiple comparisons test after one-way ANOVA. Data are presented as mean ± SEM. **n)** Normalized distribution of navigation strategies implemented by WT (left) and ChR2 (right) mice across training days in the Barnes maze. **o)** Bar and scatter plot comparing the percentage of trials in which each mouse used random (left), serial (middle), and direct (right) navigation strategies on training days 1 and 4 in non-light-stimulated WT and ChR2 mice. Each point represents a single animal. Data in the bar plots are presented as mean ± SEM. ***: P < 0.001; *: P < 0.01; *: P < 0.05; Sidak’s multiple comparisons test after two-way ANOVA.

**Supplementary figure 6.**
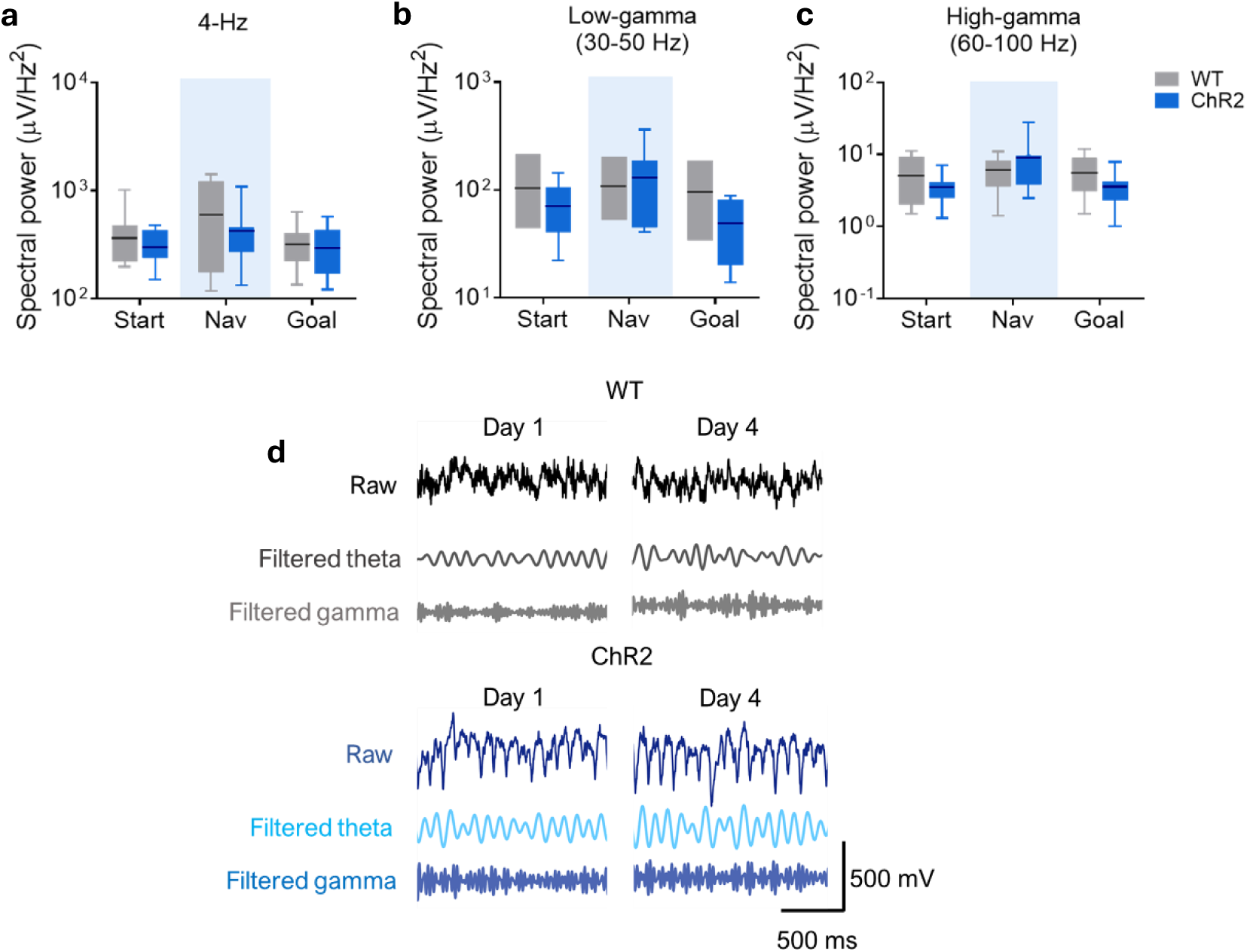
**a-c)** Box plot comparing the spectral power of 4-Hz (**a**), low-gamma (**b**), and high-gamma (**c**) oscillations in the mPFC between WT and ChR2 mice across start, navigation, and goal stages. The middle, bottom, and top lines of the box plot correspond to the median, lower, and upper quartiles, and the edges of the lower and upper whiskers correspond to the 5th and 95th percentiles. **d)** Representative raw and theta and low-gamma filtered recordings form the mPFC of WT (upper) and ChR2 (lower) mice during the navigation stage of training in the Barnes maze at day 1 (left) and day 4 (right).

**Supplementary figure 7.**
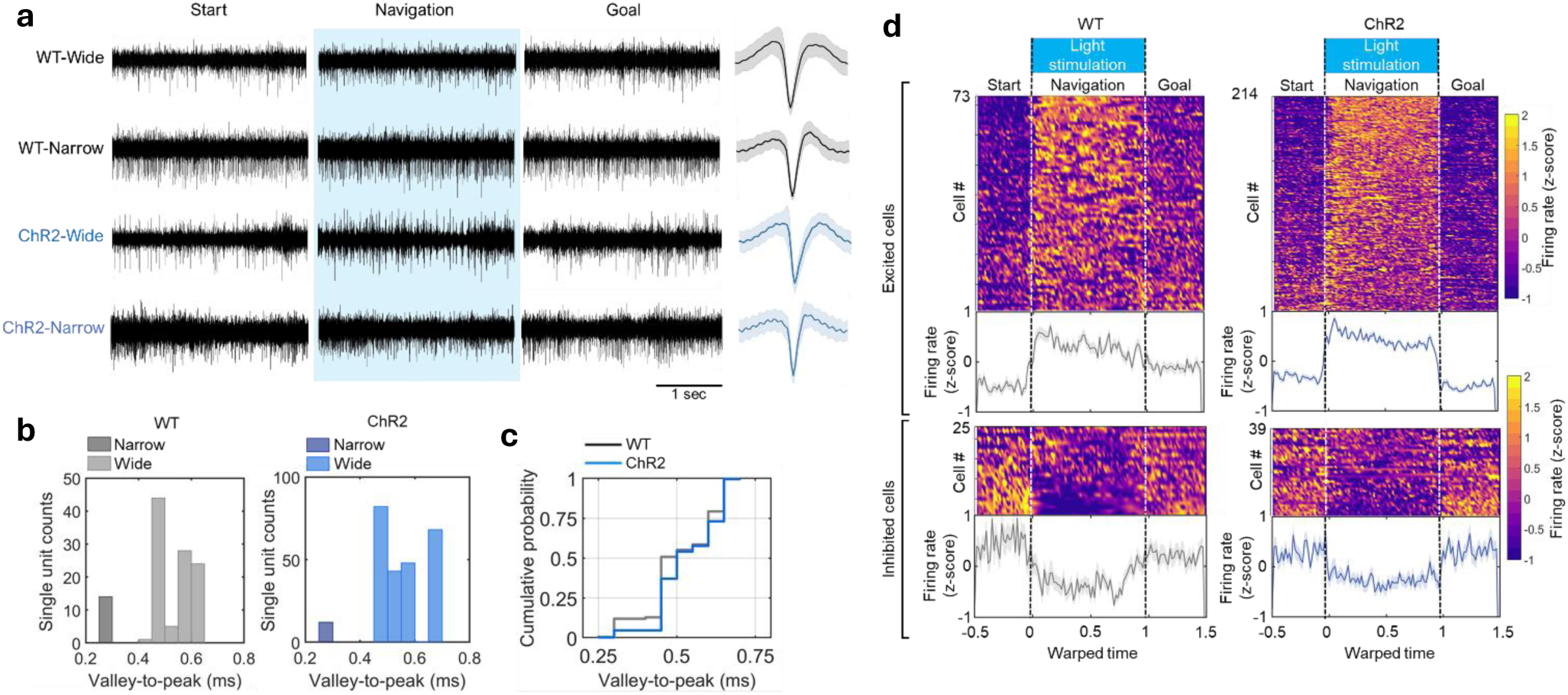
**a)** Examples of wide and narrow spike waveform of single units recorded simultaneously from the mPFC of WT and ChR2 mice during training in the Barnes maze. Each row corresponds to the same single unit recorded at different stages (start, navigation, goal) in a single session. The average spike waveform of each single unit is depicted on the right. **b)** Distribution histogram showing the counts of identified single units recorded in the mPFC of WT (left) and ChR2 mice (right) according to the valley-to-peak (VTP) duration. **c)** Cumulative histogram comparing the distribution of VTP durations of all single units recorded from the mPFC of WT and ChR2 mice. **d)** Color-coded peri-event time histogram of the normalized firing rate (z-scored) with respect to warped session time for all prefrontal single units recorded in WT (left panel) and ChR2 mice (right panel) that increased firing (upper) and decreased (lower) firing during the navigation stage of the Barnes maze. Start, navigation, and goal stages are indicated, as well as light stimulation. Each row represents a different neuron. Plots at the bottom represent the corresponding mean normalized firing rate. Shaded areas represent SEM.

**Supplementary figure 8.**
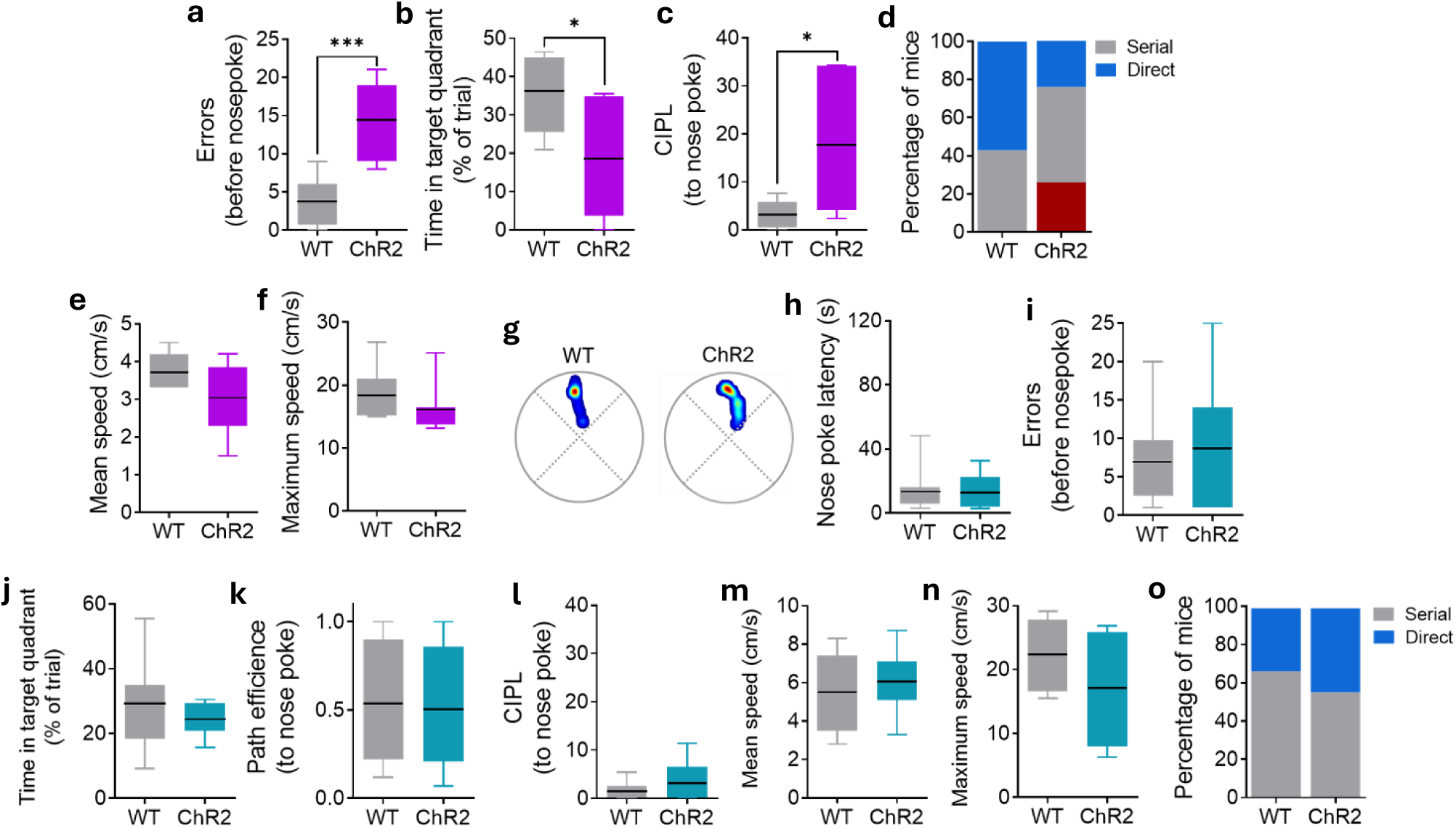
**a-c)** Box plot comparing errors before escape nose-poke (**a**), time in target quadrant (**b**), and CIPL to escape nose-poke during the memory retrieval sessions. ***: P < 0.001; *: P < 0.05; Tukey’s multiple comparisons test after one-way ANOVA. **d)** Normalized distribution of navigation strategies implemented by WT (left) and ChR2 (right) mice during the memory retrieval session. **e, f**) Box plot comparing mean (**e**) and maximum speed (**f**) between WT and ChR2 mice during the memory retrieval session. **g)** Example of color-coded occupancy plots in the Barnes maze during the memory retrieval session of non-light-stimulated WT (left) and ChR2 mice (right). **h-n)** Box plot comparing the escape nose-poke latency (**h**), errors to escape nose-poke (**i**), time in target quadrant (**j**), CIPL (**k**) and path efficiency to escape nose-poke (**l**), and mean speed (**m**) and maximum speed (**n**) during the memory retrieval session of non-light stimulated WT and ChR2 mice. For all the boxplots, the middle, bottom, and top lines correspond to the median, lower, and upper quartiles, and the edges of the lower and upper whiskers correspond to the 5th and 95th percentiles. **o)** Normalized distribution of navigation strategies implemented by non-light stimulated WT (left) and ChR2 (right) mice during the memory retrieval session.

**Supplementary figure 9.**
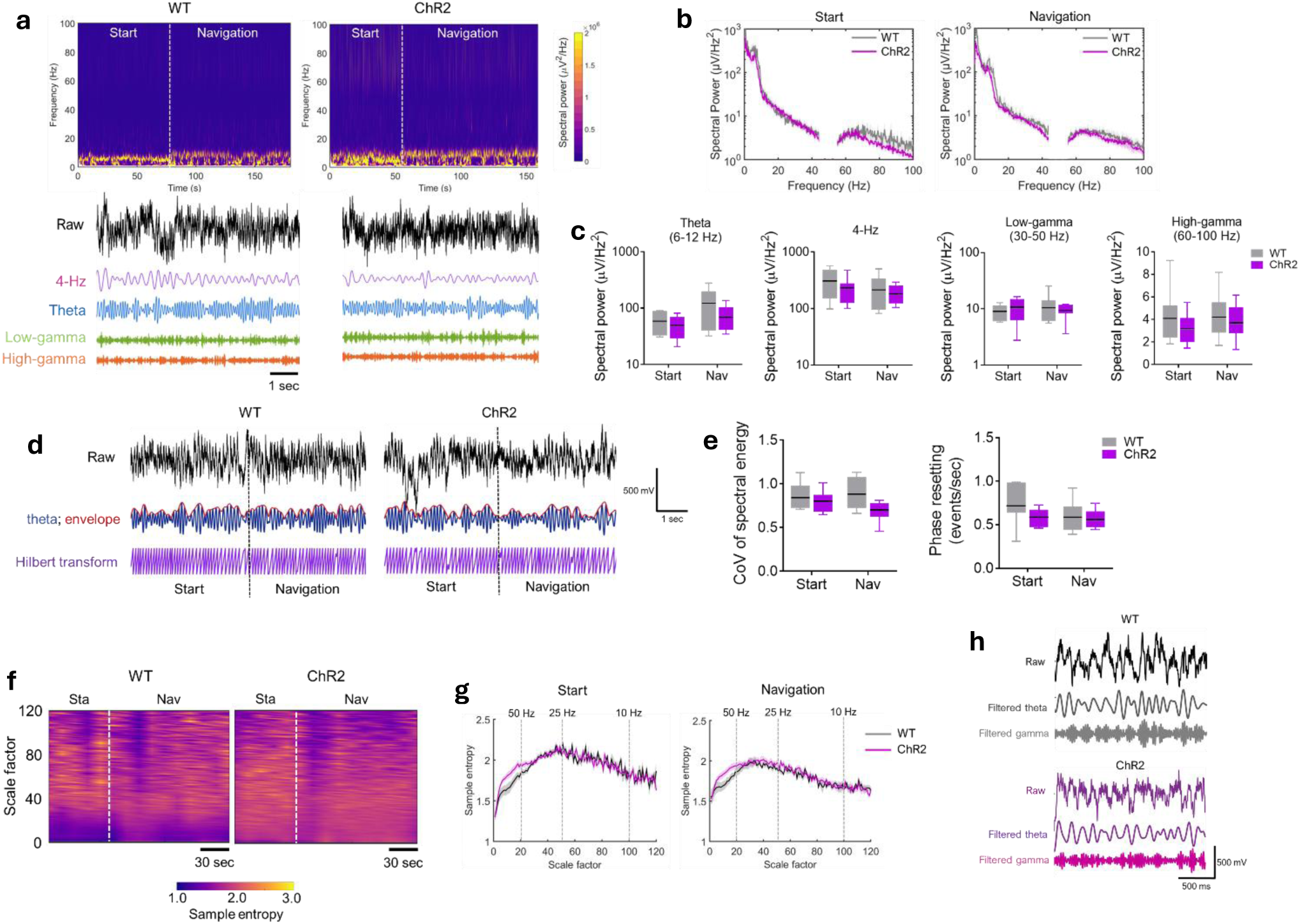
**a)** Example of time-frequency color-coded analysis of spectral power (upper panel) and raw and filtered (4-Hz, theta, low-gamma, and high-gamma) LFP recordings (lower panel) from the mPFC of WT (left) and ChR2 (right) mice during memory retrieval. Start and navigation stages are indicated. **b)** Average power spectral density analysis of the LFP recorded from the mPFC in WT and ChR2 mice during the start and navigation stages in the memory retrieval session. Data are presented as mean (solid line) ± SEM (shaded area). **c)** Box plot comparing the spectral power of theta, 4-Hz, low-gamma, and high-gamma oscillations in the mPFC in the start and navigation stages between WT and ChR2 mice during memory retrieval. **d)** Example of raw (black) and theta-filtered (blue), and its envelope (red) of LFP recording from the mPFC of WT (upper) and ChR2 mice (lower) in the transition from start to navigation stages during memory retrieval. **e)** Box plot comparing the CoV of spectral energy (left) and incidence of phase resetting (right) of theta oscillations in the mPFC in the start and navigation stages between WT and ChR2 mice during memory retrieval. The middle, bottom, and top lines of the box plot correspond to the median, lower, and upper quartiles, and the edges of the lower and upper whiskers correspond to the 5th and 95th percentiles. **f)** Example of time-scale analysis of multiscale entropy (MSE) of an LFP signal recorded from the mPFC of WT (left) and ChR2 (right) mice during a memory retrieval session. **g)** Comparison of the average MSE of the LFP signals recorded from the mPFC of WT and ChR2 mice in the start and navigation stages during memory retrieval. Data are presented as mean (solid line) ± SEM (shaded area). **h)** Representative raw, theta, and low-gamma filtered recordings form the mPFC of WT (upper) and ChR2 (lower) mice during the navigation stage of a memory retrieval session. For all the boxplots, the middle, bottom, and top lines correspond to the median, lower, and upper quartiles, and the edges of the lower and upper whiskers correspond to the 5th and 95th percentiles.

**Supplementary figure 10.**
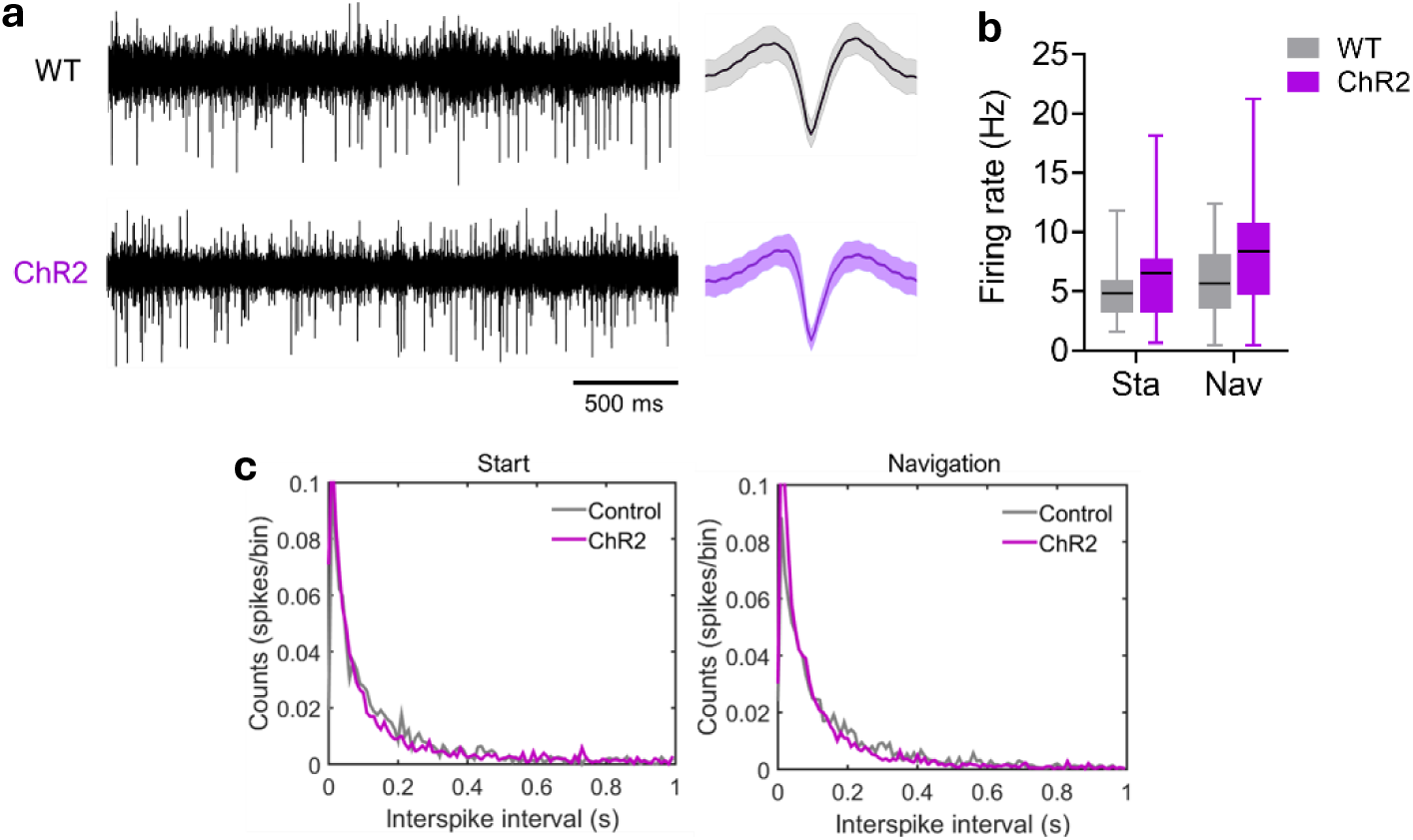
**a)** Representative single unit (LFP filtered at 500-5000 Hz) traces recorded from the mPFC of WT and ChR2 mice during the navigation stage in a memory retrieval session. The average waveform for each single unit is shown on the right. **b)** Box plot comparing the firing rate in start and navigation stages in the mPFC of WT and ChR2 mice during memory retrieval. The middle, bottom, and top lines of the box plot correspond to the median, lower, and upper quartiles, and the edges of the lower and upper whiskers correspond to the 5th and 95th percentiles. **c)** Comparison of the average inter-spike interval (ISI) between single units recorded from the mPFC of WT and ChR2 in start and navigation stages during memory retrieval.

